# Hidden Aspects of the Research-ADOS are Bound to Affect Autism Science

**DOI:** 10.1101/717827

**Authors:** Elizabeth B Torres, Richa Rai, Sejal Mistry, Brenda Gupta

**Affiliations:** Rutgers University, Psychology Dept., Piscataway, NJ; Computer Science, Center for Biomedical Imagining and Modeling; Rutgers University Center for Cognitive Science; University of Utah Medical School, Salt Lake City, UT; Montclair State University, Montclair, NJ

**Keywords:** metric space, norm, change, trajectory, probability distributions, non-parametric statistics, distribution free analyses, normal distribution

## Abstract

The research-grade ADOS is a broadly used instrument that informs and steers much of the science of Autism. Despite its broad use, little is known about the empirical variability inherently present in the scores of the ADOS scale, or their appropriateness to define change, to repeatedly use this test to characterize neurodevelopmental trajectories. Here we examine the empirical distributions of research-grade ADOS scores from 1,324 records in a cross-section of the population comprising participants with autism between 5-65 years of age. We find that these empirical distributions violate the theoretical requirements of normality and homogeneous variance, essential for independence between bias and sensitivity. Further, we assess a subset of 52 typical controls vs. those with autism and find lack of proper elements to characterize neurodevelopmental trajectories in a coping nervous system changing at non-uniform, non-linear rates. Lastly, longitudinally repeating the assessments over 4 visits in a subset of the participants with autism for whom verbal criteria kept the same appropriate ADOS modules over the timespan of the 4 visits, reveals that switching the clinician, changes the cutoff scores, and consequently, influences the diagnosis, despite maintaining fidelity in the same test’s modules, room conditions and tasks’ fluidity per visit. Given the changes in probability distribution shape and dispersion of these ADOS scores, the lack of appropriate metric spaces, and the impact that these elements have on sensitivity-bias co-dependencies, and on longitudinal tracking of autism, we invite a discussion on the use of this test for scientific purposes.

## 1. Introduction

Several debates have been published surrounding controversial reliance on the research-grade Autism Diagnostic Observational Schedule (ADOS) [1] for use in scientific Autism research [2]. The field seems to have reached a point whereby some Autism researchers advocate the importance of nervous systems’ biorhythms [3; 4; 5; 6; 7; 8; 9; 10] and the use of biophysical data to adapt tenets of Precision Medicine [11; 12] to a nascent field of Precision Psychiatry [4; 13]. These new developments have led to a surge in the interest to develop digital biomarkers for personalized approaches to diagnose and treat disorders of the nervous systems [14]. Among these, are neurodevelopmental disorders whereby the child’s nervous systems develop differently, and maturational milestones are not met at the typically expected times [6] (Figure 1). Often, these different courses of maturation give rise to somatic-sensory-motor problems that result in highly heterogeneous manifestations. Such manifestations constitute observable symptoms useful to build categories that could provide criteria for a diagnosis [11; 15; 16] (e.g. as informed by those used in the Diagnostics Statistical Manual DSM-5 from the American Psychiatric Association APA [17]).

**Figure 1.**
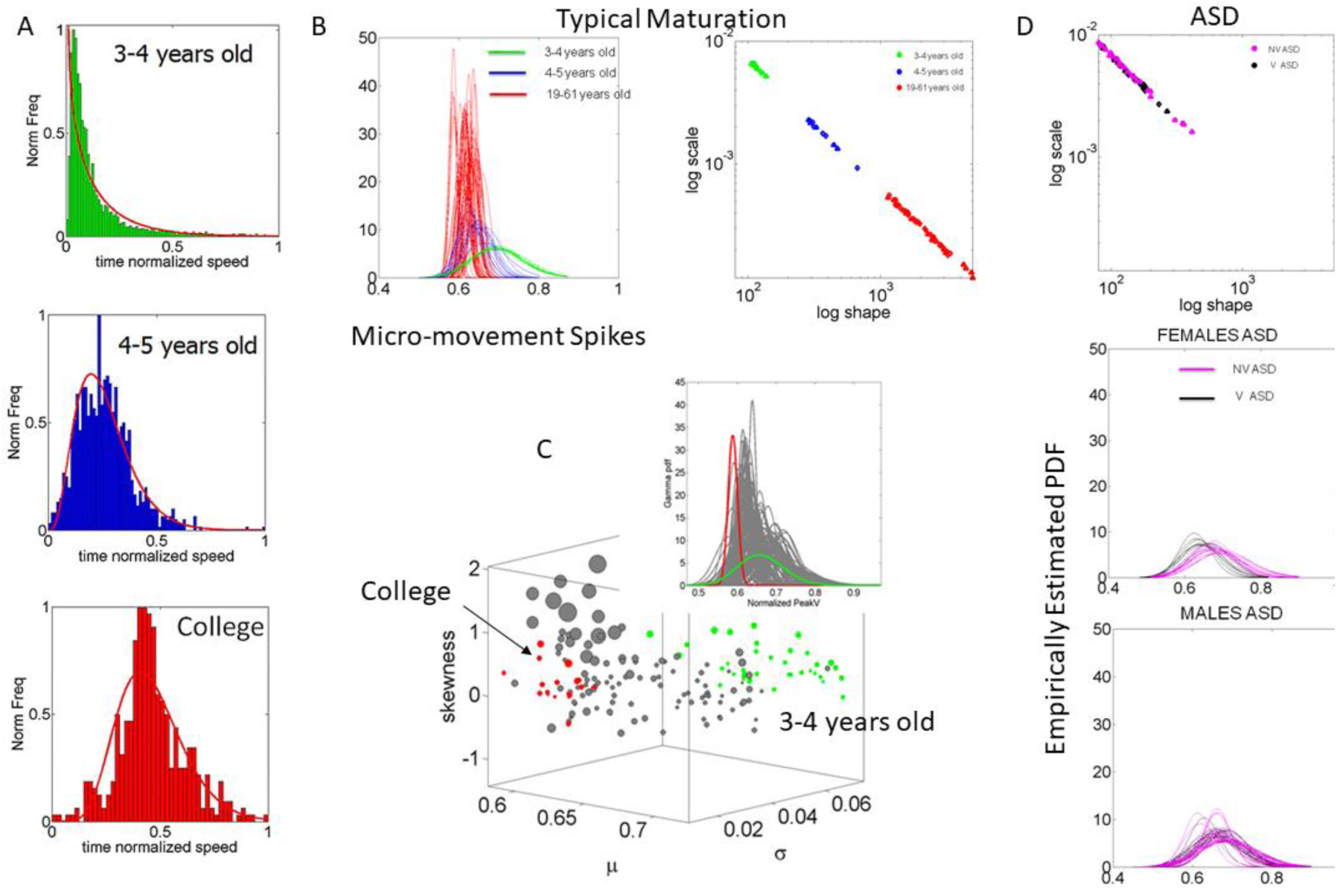
Digital biomarkers at work in Autism. (A) Typical maturational trajectory of unaffected individuals characterizing the somatic-sensory-motor variability of pointing speed over the span of 30 years, taken cross-sectionally in a random draw of the population. Frequency histograms show the lack of normality in motor variability in 3-4 years of age. By 4-5 years of age, the Exponential distribution shifts and a heavy tail appears indicating faster motions and lower noise-to-signal ratio in speed variations evolve as well. Individuals of college age begin to show more symmetric distributions with high signal content and predictive patterns more accurately forecasting the sensory consequences of their impending actions and decisions. (B) A characterization of data in (A) under a standardized data type using a Gamma process to empirically estimate the shape and scale parameters best characterizing the empirical distribution in a maximum likelihood estimation sense. The normalized data provide the probability density functions PDFs and the Gamma parameter plane shows the course of maturation of (A) whereby each dot represents a participant (personalized approach) spanning a family of Gamma PDFs. This maturation follows a power law across ages of the population whereby as the distribution becomes more symmetric (to the right of the shape axis of the Gamma parameter plane), the noise-to-signal ratio decreases along the scale axis. Matured systems land in a low noise and high symmetric distribution regimes, whereas immature systems remain in a random (Exponential) and noisy regime where events in the past do not contribute to the forecasting of future events. (C) The stochastic signatures of the Gamma moments separate extreme states of typical maturational stages defining boundary values of normative stochastic data. (D) The prevalence of high noise and lack of predictive code is evidenced in Autism from 3 to 30 years of age. The speed variations show patterns of random noise and remain as those of a typically developing 3-year-old child, whether the Autistic person is verbal or non-verbal, male or female. Digital biomarkers derived from motor-sensing variability identify target for treatments (dampen the noise) and provide a natural classifier of lifelong nervous systems disorders. (Pictures previously published in [6; 14; 18]).

Perhaps part of the problem stems from the rather broad umbrella term that clinical criteria produces, encompassing several comorbid conditions, and thus leading to highly heterogeneous disorders. In many fields, the clinical criteria to categorize neuropsychiatric and neurological symptoms, tend to shift over time, often informed by scientific progress. In Autism spectrum disorders (ASD) however, upkeeping with the science has proven rather challenging. This is because basic research and clinical inventories are intertwined. Unlike in other fields, according to AI-based analyses of over 17,000 Autism-related peer-reviewed publications in the US, spanning since 1994-2015, clinical criteria tend to drive Autism research [19]. This occurs under a “*one size fits all model*” that obfuscates progress, partly due to complete absence of characterizations of typical development to provide proper metrics defining normative states of social and emotional developmental trajectories.

The ADOS test is not a norm-reference test. It is a criterion-reference test. As such, it should not be adopted to inform scientific research about longitudinal change. Because this is not a metric test, it is not possible to derive a similarity metric to quantify change and its rate, so important in developmental sciences. Despite these issues, the ADOS scores are used to assess longitudinal developmental trajectories [20; 21] under assumptions of normality and while adopting linear scales to examine the data. We note here that a trajectory requires a norm to measure the length of the velocity vector comprising change of position (of a parameter) over time. In the absence of a metric (to measure length) and in the presence of a rather non-linear complex dynamical system growing and changing by the day in a non-linear accelerated manner [4], such use of the ADOS test poses a challenge to Autism science.

Indeed, inherent in the use of the Autism clinical model that has been transferred to basic science as a research-appropriate model, are the assumptions of normal distribution and linear processes with stationary statistics that emerge-independent of the observer’s bias when rating rather complex and dynamic social behaviors (Figure 2AB). In research adopting such scoring scales, the scales have yet to be mapped to biophysical data from the somatic-sensory-motor neurophysiology underlying the types of biorhythmic motions that make up all behaviors. The word “*behaviors*” in the clinical field of Autism has in fact a very different meaning than it does in basic research (e.g. in the field of Behavioral Neuroscience), triggering a negative connotation that makes it a rather controversial term [3]. Nevertheless, the biophysical data underlying the types of observed behavior that lead to a diagnosis of Autism, produce empirical outcomes that may not correspond to the theoretical statistical assumptions underlying the analyses that defines the clinical scale used in basic research (Figure 2CD and inset in D). The lack of normality in the distributions of various biophysical parameters of basic electrophysiology appears across several lines of enquiry, spanning from imaging to behavioral physiology [5; 14; 15], but it has not been investigated in research-grade ADOS scores empirically obtained from the adopters of the test in Autism science.

**Figure 2.**
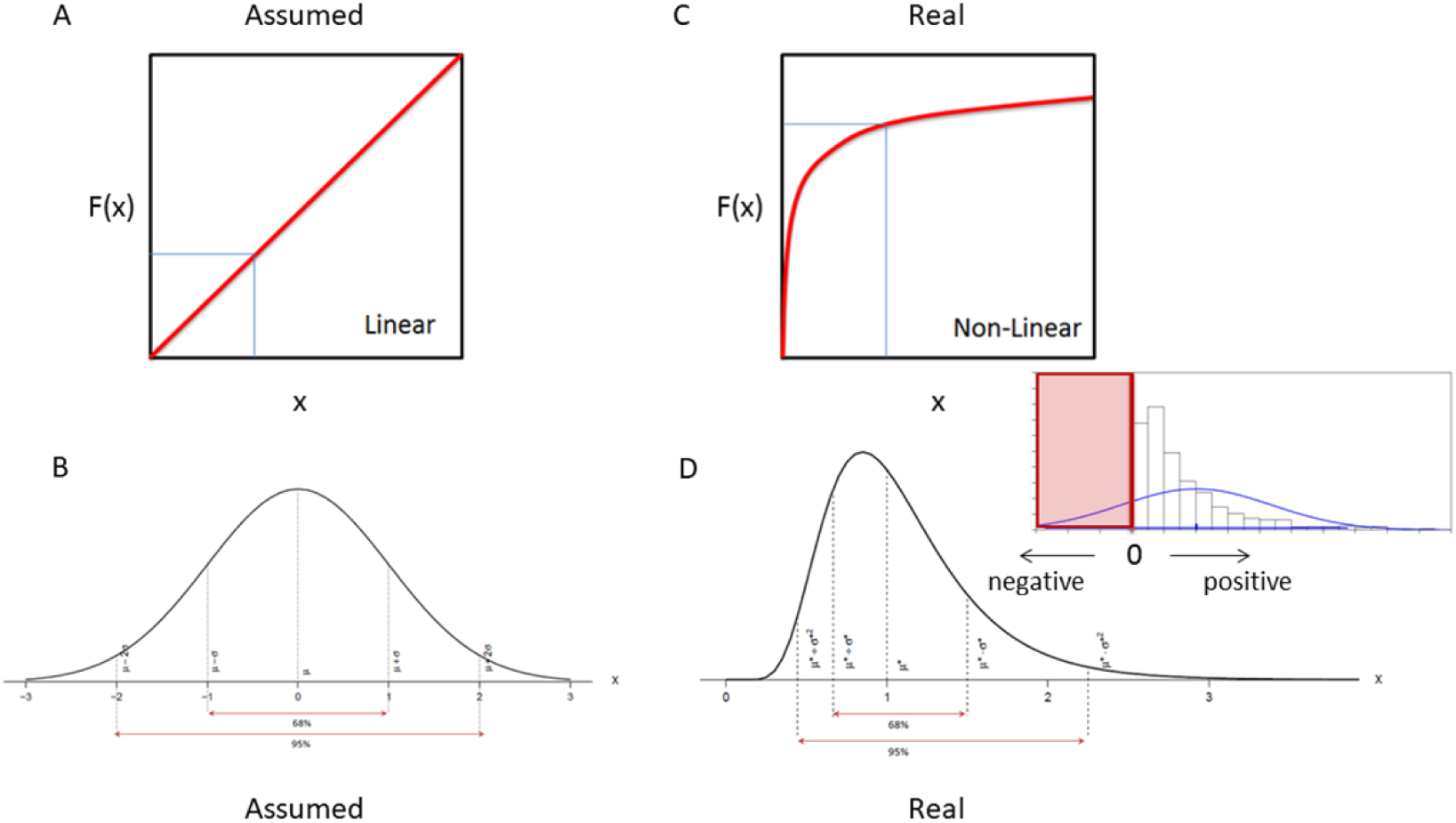
Assumed theoretical models vs. empirical reality. (A) Linear models are routinely used to characterize neurodevelopment under a “*one size fits all*” approach (B) that assumes normality in the neurodevelopmental variability and focuses on a theoretical mean at the expense of smoothing out as noise important fluctuations in the data. (C) Neurodevelopment occurs along non-linear accelerated trajectories that span non-uniform distributions of various kinds. (D) In neurodevelopmental data, the presence of non-Gaussian distribution is ubiquitous across different aspects of neurobiology, from molecules to behavior, to basic physical growth. Often, the parameters under investigation span positive values with 0-value at the lowest bound. For example, the speed of the hand during pointing behavior is 0 at rest and increases away from the body toward the target with limiting upper bound depending on the length of the person’s limb and the level of elasticity of the arm’s joints, the level of muscle force, among other factors. Such distributions tend to be skewed and when a symmetric (theoretical) distribution is imposed on the data, there is the danger of producing spurious values in the negative range, e.g. when expressing error bars with +/-two standard deviations from the often-assumed Gaussian mean.

In this work, we ask if the theoretical statistical assumptions made by the adopters of the research-grade ADOS used to characterize Autism in basic science, correspond to those derived from the empirical data that the use of such test produces. We have reasoned that the sensitivity and reliability tests that researchers adopting this clinical scoring system assume to assess its validity, are built under very specific statistical assumptions of normality [22; 23; 24; 25]. Under such assumptions, the shape and dispersion of the variability from the scores’ probability distributions are critical to maintain independence between the inherent bias of the observer and the sensitivity of the test.

The ADOS range of scores is based on positive integer values, with 0-value at the lower bound, representing that the behavior is not present as specified. Behaviors coded on the ADOS are assumed to be behaviors that occur in non-spectrum individuals (e.g., eye contact, pointing, shared enjoyment) as well as behaviors that could occur in ASD (e.g., stereotyped/idiosyncratic language, complex mannerisms). We note that those behaviors present in the non-autistic individuals have not been experimentally assessed under ADOS conditions, as noted by their classical paper [1], in the last sentence of the paper. Aspects of social interactions such as eye contact, pointing movements, face-processing micro-motions, etc.) are routinely studied in the neuromotor control developmental literature using objective means that characterizes several types of motions (e.g. [6; 9; 10; 26; 27; 28; 29]), but that literature has not crossed paths with the ADOS.

It may be possible that the *discrete* data from the observational scores statistically aligns well with the non-normally distributed, non-linear complex dynamics data that we find in the *continuous* biophysical data underlying the physiological and neurobiological studies of Autism. If that was the case, we would have to rethink the scales currently defining the cut-off criteria adopted in research to detect Autism, as such discrete (static) scores are often expected to correlate with the continuous (dynamic) biophysical data from scientific studies.

We may also need to begin to think about new mathematically appropriate ways to track neurodevelopment while using behaviorally based observational means. For example, in pediatrics, the Growth Charts from the CDC and the WHO serve the purpose of establishing normative criteria to measure departure from typical development, as the child physically grows. In Autism, there is nothing comparable to the growth charts, so there are no metrics of similarity to detect and track *change* and its rate. It has now been established that observational behavioral criteria grounded on psychological (subjective) constructs drive the scientific quest in Autism and supersedes physiological (objective) criteria [14; 19], so the time is ripe to build a similarity metric based on physiological data registered from the fast growing and rapid developing nervous systems.

One fundamental difference between observation-driven clinical diagnostics tools enforced in Autism research, and the physiological data that we have gathered across labs to address basic scientific research in Autism (e.g. to characterize genetics, brain activity and somatic-sensory-motor issues across different imaging and wearable sensors platforms, e.g. [5; 6; 30; 31; 32; 33; 34]) is that there is always a group of neurotypical controls included in the study. This is so, to help us establish normative data and to ascertain departure from it across the human lifespan. In contrast to the Autism research involving physiology and more generally neurobiology, we could not find any study characterizing the neurotypical population using the research-grade ADOS defining criteria for Autism (i.e. based on social interactions and communication, and ritualistic repetitive behaviors, RRBs.) This is so because the clinical version of the ADOS is not a norm-referenced test, but rather a criterion-referenced one.

As such, the research-grade ADOS which was adopted in science from the clinical counterpart, has no proper *metric* to measure relative changes away from typical levels. In this sense, although the ADOS modules were designed to account for possible disparities in cognitive/verbal capacity, there is no age-dependent criteria in the research-adopted version to ascertain physical rate of change and to measure departure from typical physiologic maturation, i.e. maturation with respect to normative trends (Figure 1A). Even the so-called “standardized” ADOS severity scores do not address this point, because as they were developed for criteria-referenced for clinical use, the scale was not built using typical controls as a norm-referenced test would be [35; 36; 37]. The score is designed to compare an individual with ASD to other individuals with ASD of the same age and language level. It also has a range of 6-10 for individuals with ASD and is not meant to represent ASD on a range of 1-10. Autism is not only highly heterogeneous. It also has neurodevelopmental asynchronies in a group of the same age, meaning that two individuals may be 10 years old, but one may have the signatures of neuromotor control from normative 3-year-old children [6] (Figure 1A, B, D). Thus, aging with autism is different than typically aging.

The absence of a *similarity metric* for the adopted research ADOS poses a challenge to the scientific community. What is the normative range of scores that reflect age-appropriate typical social interactions?

Because of the prevalent influence that the research-grade ADOS test has on basic science at all levels, it is imperative to examine the inherent theoretical assumptions that adopters of this test have made and verify that the outcome of this test -as administered in research settings-empirically matches the theoretical assumptions of the users.

In the first part of this paper, we use the Autism Brain Imaging Data Exchange (ABIDE) repository [38] to examine *1,324* clinical records of participants with ASD, with the purpose of better understanding the inherent statistics of the ADOS scores obtained in research settings. Specifically, we ask if the current theoretical statistical assumptions that the adopters of the research-grade ADOS inventory make are in correspondence with the empirical cross-sectional (publicly available) data from the Autistic population across the human lifespan (*5-65* years of age) (Figure 3). In the second part of the paper, we examine the ADOS-2 scores’ distributions in 52 participants, 26 typical controls with no health or mental issues, and 26 participants suspected to have (and confirmed to have) Autism. We ask if the neurotypical participants cluster at values significantly away from the 0-value denoting the absence of a behavior that would be otherwise present; or if they significantly depart from 0. In the last part of the paper, we longitudinally track a subset of the individuals with Autism who returned to the lab 4 visits. We tested these participants using the same module (twice), each module administered by different clinicians, to assess the extent to which such changes impacted the scores that they received, despite adjusting for physical age. We report our findings and encourage new transformative changes towards objective behavioral characterizations to help advance Autism science toward the use of similarity metrics that appropriately measure change and its rate along properly quantified neurodevelopmental trajectories.

**Figure 3.**
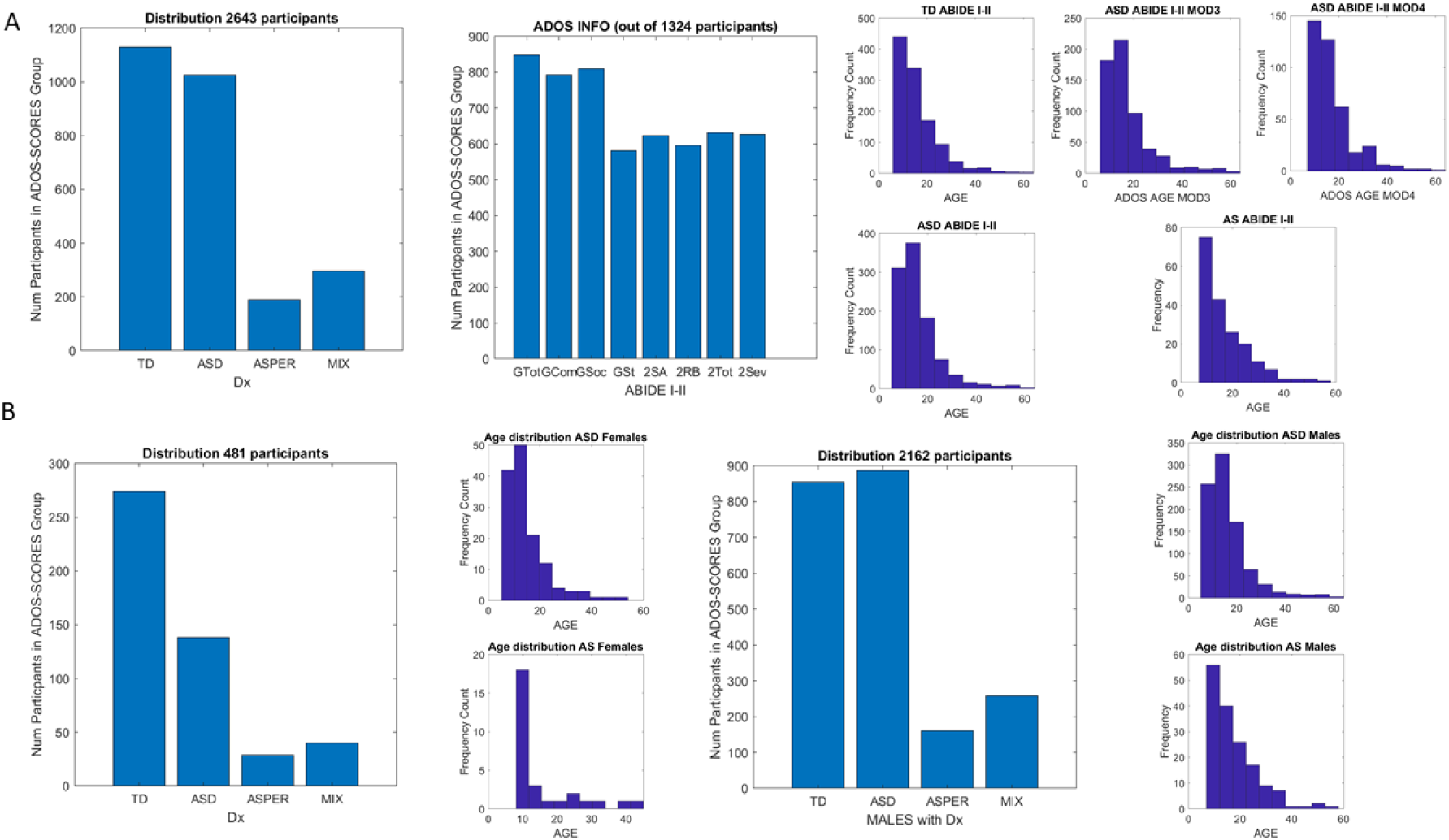
ABIDE data sample. (A) Participants (2,643) according to the diagnosis and ADOS information from 1,324 participants in the spectrum of Autism divided by sex 481 female vs 2,162 male participants (B). Histograms are also divided by age and sex for TD, ASD, AS and distributions of participants who performed Module 3, or those who performed Module 4.

## 2. Methods

The Rutgers University Institutional Review Board (IRB) committee approved all protocols used in this study. Parental and /or legal guardian consent was obtained for all participants. All procedures were performed in compliance with the Helsinki Act under IRB approval. For the consent process, we read and explained to the parents the IRB-approved protocol and the child assent form. After the informed consent process, all 26 families agreed to participate and signed written parent permission and assent forms. The participant’s name and identifiers were removed to maintain confidentiality across the entire analyses. Further, deidentified data were used in all sections of the manuscript, including the **Tables**.

### 2.1. The ADOS Assessment

The Autism Diagnostic Observation Schedule assessment is a tool to aid the diagnosis of autism. It is often used in combination with other tools such as the Diagnostics Statistical Manual DSM-5 and the ADI-R screening tool. ADOS had at some point four different modules (now 5 modules and training for toddler’s and adult’s levels are more common.) Each module is designed to provide the most appropriate test for an individual at a certain language level. There is a newly created calibrated severity score (CSS) in ADOS-2. It is based on the person’s age as it converts an individual’s Total ADOS-2 score in comparison to other individuals with ASD at the same age and language level (for each individual module). For the actual ADOS-2 administration age does not determine the module but may determine algorithm items within that module. We note however, that two children of the same age will very likely have very different neuromotor control age, as assessed by objective physiological metrics. In general, age has no real meaning in neurodevelopmental disorders, where the coping nervous systems of different individuals born at similar dates, evolve at very different rates [6; 39] and see Figure 1.

In this experimental set up, the most appropriate module was determined by two experienced clinicians, a Developmental Pediatrician and a Developmental Clinical Psychologist. Further, two clinically certified raters independently video-taped and discussed the sessions to ensure module administration fidelity.

The two raters who administered the ADOS to the participants were research reliable. Research reliable examiners are trained professionals who have undergone ADOS-2 Clinical training. ADOS-2 Clinical Training is an introductory workshop with instructional methods that include lectures, videos, demonstrations of administration and scoring, and discussions. This workshop serves as a prerequisite for more thorough training required to obtain research reliability needed to use the ADOS-2 for research purposes.

Research reliable means that perspective trainers have submitted ADOS-2 administration videos to fellow research reliable examiners and have received at least an agreement of 80% of similar scores on the ADOS-2 administration as displayed through the submitted video. The perspective research reliable trainee and the already research reliable trainer have at least 80% of similar scoring on the ADOS-2 administration video submitted by the trainee. Three videos must be submitted and a minimum of 80% similar scores must be achieved. Once they passed the requirement, they became research reliable ADOS raters. The information about four of the Modules that we used are taken from the manual and briefly summarized below but are not meant to be complete (see also Supplementary table) and when in doubt, consult the ADOS manual:

#### Module One

It is designed for individuals who do not have consistent verbal communication skills and the tasks use completely non-verbal scenarios for scoring.

#### Module Two

It is designed for individuals who have minimal verbal communication skills, including young children at age-appropriate skill levels, whereby tasks require moving around the room and interacting with objects. All objects are standard and come in a standardized kit.

#### Module Three

It is designed for individuals who are verbally fluent. These participants are also capable of playing with age-appropriate toys. The test is conducted largely at a desk or table. In our setup, the table was always the same and it was positioned in the room in the exact same configuration. The room where the test took place was not changed from session to session and participant to participant. Researchers and ADOS-certified personnel in the study (4 total) made sure that the conditions were identical in each session, task, child, rater and module.

#### Module Four

It is designed for individuals who are verbally fluent but no longer at an age to play with toys. This module incorporates some Module Three elements, yet it is more conversational regarding daily living experiences.

Often the examiner may choose one module and realize that the participant’s functional abilities anticipated by that module do not match the rater’s expectation. Then the tester may choose another module. This is a common practice. As such, the present experiment manipulated the module type, to probe the participant’s responses to the same module administered by two different raters, or to determine the adequacy of the modules.

The modules involving playing with toys or objects have the tester present standardized scenarios to evoke responses and rate the child’s performance. Some elements of the game (e.g. a puzzle) are left out on purpose to evoke the need in the child to ask for other pieces. The examiner uses several strategies to evoke different responses and assess the child’s reaction. This mode of testing behaviors inherently makes certain assumptions about social expectations that do not necessarily transfer from culture to culture. Further the ADOS states that no sensory-motor issues are present without ascertaining the intactness of the child’s peripheral nervous systems before administering the test. A child response may not be voluntary. When the nervous systems are damaged, there is an inevitable component of the response that is never evaluated during the test, owing to the test’s assumption that no significant sensory and motor issues are present. As such, it is never possible to assess causality. For all these reasons, and because our lab receives a highly diverse population from many different cultures and neuromotor developmental stages, the experimental protocol manipulated the variable representing the rater, while maintaining constant all other conditions (context, module, room, lay out of all objects in the room, etc.) and videotaping the sessions by other two independent ADOS-certified raters. As it is required from the ADOS, raters flexibly adapted each session to the child’s responses to the flow of the tasks, while maintaining fidelity to the tasks employed in each module of choice.

The response of the child determines the score. Likewise, the way in which the rater evokes the response influences the child’s choice of actions that are consequential to the rater’s provoking actions. To probe the extent to which a change in the tester influenced the scoring, our experiment manipulated the rater as a parameter, while holding all other conditions constant in two different visits. Participants came to the lab a total of four visits. For each participant, two modules were selected, and research-reliable testers were employed. One module was rendered the most adequate one, while the other was rendered the feasible one. By most adequate, we mean that the module was at the child’s verbal level and developmental level, while feasible means that the child could perform the entire module, but it would not be the adequate one to perform a diagnosis or to aid a clinician in performing a diagnosis of autism. We note that previous research indicates that inappropriate ADOS module use invalidates the assessment and the scores do not accurately reflect the child’s performance on the assessment. Nevertheless, since this study is not about diagnosing autism, but rather it is about evaluating the use of this ADOS test in basic science, specifically assessing the variability that using different raters may add to the scores; in addition to changing the rater, we are also manipulating the use of the modules across different visits.

Each module took between 40-60 minutes to complete. Both the rater and the participant were recorded by two video cameras from different angles and by wearable smart sensors that they wore embedded in the clothing, as wrist watches on the wrists and on the ankles. The digital data will be the subject of a different paper. Additional information about the ADOS test can be found in the Supplementary Table.

##### Open Access Data

The ABIDE records contain ADOS-2 and ADOS-G scores that we extracted to plot the distributions of these tests sub-scores across *1,324* participants, ranging between *5-65* years of age. We present the distribution of participants in Figure 3 comprising clinical records from ABIDE I and II. Preliminary results from this work involving involuntary head motions during resting state in fMRI studies were published elsewhere [5; 40; 41]. Here we focus on the nature of the distributions of the ADOS scores, assumed to be symmetric by the ADOS manual and by reliability-validity tests commonly used in clinical Psychology to validate this diagnostic tool driving basic scientific research, e.g. [42; 43].

Clinical tools for diagnoses use the tenets of Signal Detection Theory, SDT [44; 45], albeit in a black-box approach that has never verified the implicit assumptions made by the statistical packages that such papers report. Non-parametric methods to correct for non-normality are inadequate in a diagnostics test [23]. For example, in the ADOS, as in many other clinical tests, the clinician plays the dual role of being the stimulus (via the prompts to evoke social overtures and/or primed responses) and at the same time, being the observer, scoring the participant’s response. As such, the outcome of this test, which is presumed to measure social interactions between two people, heavily depends on the clinician. In the absence of the required assumptions of normality and variance homogeneity, the sensitivity (*d’*) and response bias (*β*) do not represent accurate and independent measures [23; 45]. There are co-dependencies between them.

Because of the broad use of this test in Autism research, and the possibility that the assumptions required for research validity may not be met, it becomes crucial to verify the normality assumption along with the assumption that the rater’s bias and response are independent. To that end, we take the unique opportunity that ABIDE offers with thousands of records providing the ADOS-G and ADOS-2 versions of the test for DSM-IV and DSM-5 criteria, and we examine the distributions of the ADOS scores. We gather the scores in frequency histograms built using various binning procedures [46; 47; 48]. Then we fit the normal distribution and other distributions (e.g. lognormal, exponential, Gamma, Weibull) using maximum likelihood estimation (MLE) methods in MATLAB.

We set the null hypothesis that the data comprised by the research-grade ADOS scores of the ABIDE repository came from a distribution in the normal family, against the alternative hypothesis that it does not come from such a distribution, using a Lilliefors test [49].

Besides testing the distribution of ADOS-G and ADOS-2 scores across the full set of ABIDE, we also separate the clinical data by Module 3 *vs*. Module 4. Further, we separate the data from the females and the males in the ABIDE repository-because it has been previously found that there are fundamental physiological differences between their involuntary head motion patterns derived from the images in ABIDE [41]. Typical females, females with ASD and females with Asperger’s Syndrome can be physiologically stratified and separated from the males using involuntary micro-movements [26; 41]. Further, they can be separated using voluntary movements [26]. Here we ask if the ADOS scores separate them too, or if unlike physiological criteria, the clinical ADOS scores confound males and females. All studies in the ABIDE data repository were performed under IRB approval in accordance with the Helsinki Act.

##### In-person Data Collected at the Lab

In the second part of the study, we measure the outcome of the ADOS-2 test in 52 individuals, 26 controls (ages 7-66 years old) and 26 suspected to have (and then diagnosed with) Autism spectrum disorders (4-20 years old). We used the baseline visit 1 to characterize the participants diagnosed with ASD and those who were neurotypical. This was done to ascertain the extent to which the scores’ range from typical control participants deviate from the 0-scores denoting the absence of behavior otherwise present and contributing to the overall cut-off number. We were motivated by the critical need to create a similarity metric for Autism research. Given the adoption of the research-grade ADOS for research and the fact that this observational inventory is not a norm-reference test [1], we ascertained the spread of scores obtained from neurotypicals away from 0. We quote from the ADOS-G paper: “*Replication of psychometric data with additional samples including more homogeneous non-Autistic populations and more individuals with pervasive developmental disorders who do not meet Autism criteria, establishing concurrent validity with other instruments, evaluation of whether treatment effects can be measured adequately, and determining its usefulness for clinicians are all pieces of information that will add to our understanding of its most appropriate use*.” (emphasis added). Table 1 shows the 52 participants’ scores and ages at baseline visit where the most appropriate ADOS-2 module was selected for each child.

In 14 of the individuals with ASD (mean 9.3 years old ± 3.0), we re-assessed them across 4 visits taking place within 1.3 years on average (± 6 months), to ask the extent to which switching the clinician and / or the ADOS-module would change the outcome of the test for the same child. To that end, for each child, in the first two visits, we used the same clinician but used two different ADOS-modules. According to each assessment in each visit, the raters determined the modules that were the most appropriate and feasible. From these assessments the 14 individuals described are those for which the second round of visits (visit 3 and 4) retained the same appropriate and feasible modules despite the passage of time.

The first module (visit one) determined the most appropriate module at baseline, for the given child. The second module (visit 2) was feasible (the child could do it) but was not appropriate. For example, if the most appropriate module in visit 1 was module 3, we would choose module 2 for visit 2. Then the same clinician would give these two modules whenever the participant retrained the set of modules from visits 1 and 2, according to the raters’ evaluation; and the same two modules were then administered in subsequent visits (by a different rater); module 3 in visit 3 and module 2 in visit 4 (see Table 2.)

We switched clinician and maintained module fidelity and identical room set up. In this way, each child had a chance to become familiar with the 2 modules by the time that we switched the clinician. Those two same modules in the same order as the first two visits, would then be administered by the new clinician to give us a chance to probe the influences of the clinician on the child’s response. The flexibility in task administration according to the child’s responses was respected to ensure fluid responses.

We hypothesized that this switching of clinicians (despite the use of the same modules’ and tasks’ order in each administration), would have a substantial effect on the ADOS sub-scores, thus significantly impacting the reliability of the total score and the cut-off for the diagnosis given to the child by the rater (clinician). To test this hypothesis, we used non-parametric statistics whereby we do not assume any distribution a priori. Table 2 shows the 14 scores across the 4 visits.

Lastly, since the passage of time inevitably affects any longitudinal study, and since development occurs in these children under unexpected / hidden coping mechanisms of their physical development, we also normalized by age at the time of each visit. In addition to the use of the absolute scores above (to test for size effects), we use relative scores (to test for derivative effects) assessing the rate of change, in scores per age quantity. The latter denote dynamic changes expected to occur in childhood, owing to the non-linear accelerated and irregular nature of neurodevelopment [50]. As in the previous analyses, we employ non-parametric tests.

## 3. Results

### 3.1. ADOS-G and ADOS-2 Scores do not Distribute Normally

The ADOS-G and the ADOS-2 scores for each of the criterion comprising the total, communication, social and stereotypical behaviors across ages in ADOS-G and the total, severity, social affect and repetitive ritualistic behaviors in ADOS-2 were not normally distributed. Figure 4A shows the frequency histograms taken across ABIDE I and II scores for each of the above-mentioned criteria of the ADOS-G, while Figures 4B-C break down the scores for Module 3 and Module 4 (note that ABIDE does not report Module 1 and Module 2 is very sparsely used). Figure 5 shows the corresponding frequency histograms for ADOS-G. The empirical cumulative distribution functions (CDFs) for each criterion are shown in Figure 4D for ADOS-2 and Figure 5D for ADOS-G. They were tested separately as they are different ADOS versions, as were each sub-score distribution. Notice here the similarities across CDFs for Module 3 and Module 4 and the similarity of each of these empirically estimated CDFs with the CDF corresponding to all scores. We note that pooling all scores is not valid, yet we do it here to show that the shape and dispersion of these histograms are quite similar despite the differences in Module’s tasks.

**Figure 4.**
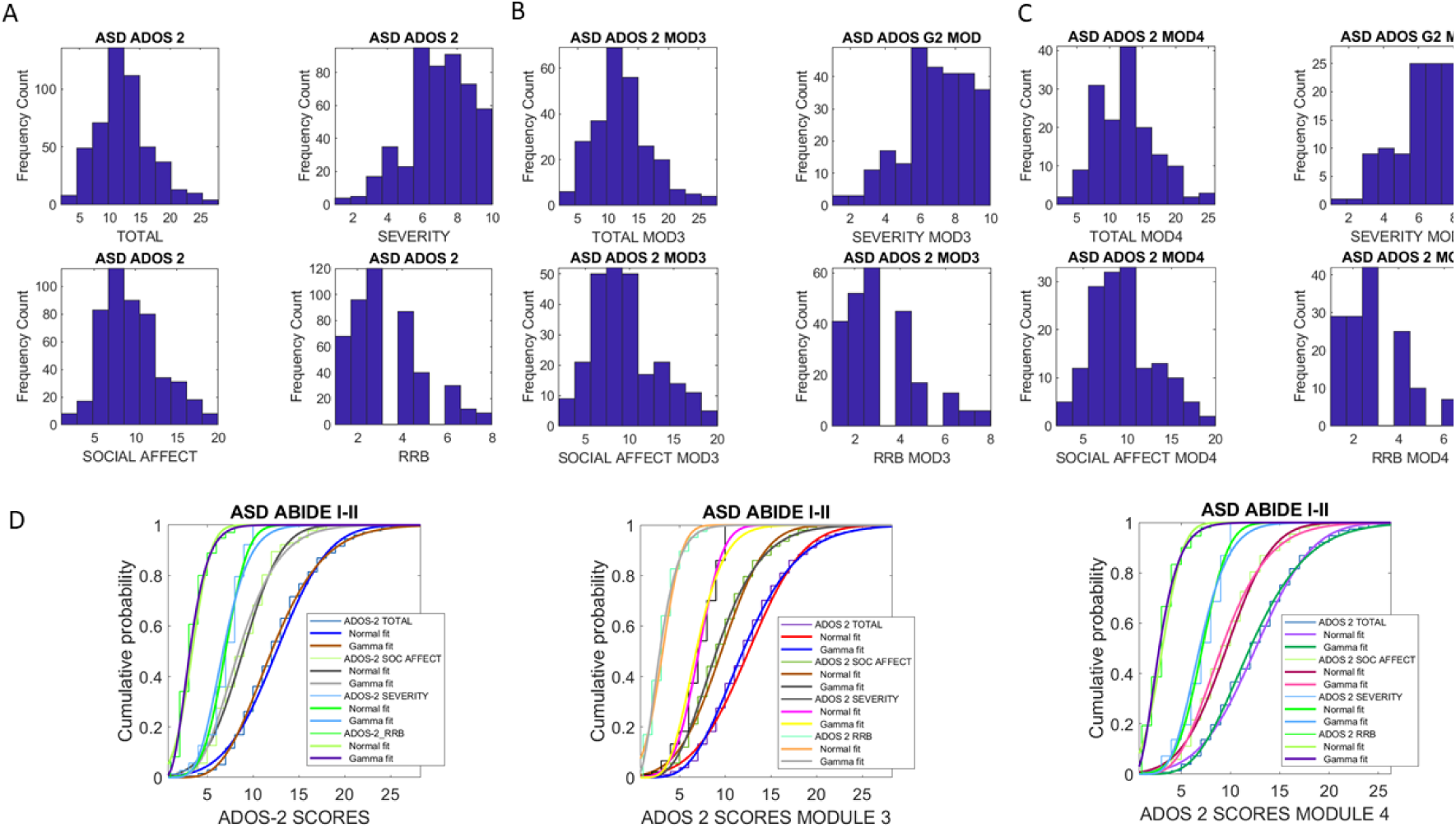
Non normality of the ADOS-2 total and each of the sub-scores distributions in the ABIDE I, II sites. (A) Empirically obtained frequency histograms for each of the ADOS-2 total and sub-scores from all ABIDE scores. (B) Same as in (A) but restricted to Module 3 scores. (C) Same as in (A) but restricted to Module 4 scores. (D) Empirically estimated CDFs for each sub-score for the Normal and Gamma distributions fit to the data. All cases failed the test of normality (see text) but no statistically significant differences were found between the empirical CDFs generated by Modules 3 and Module 4. No differences were found either between each of the Modules and the pooled data in (A).

**Figure 5.**
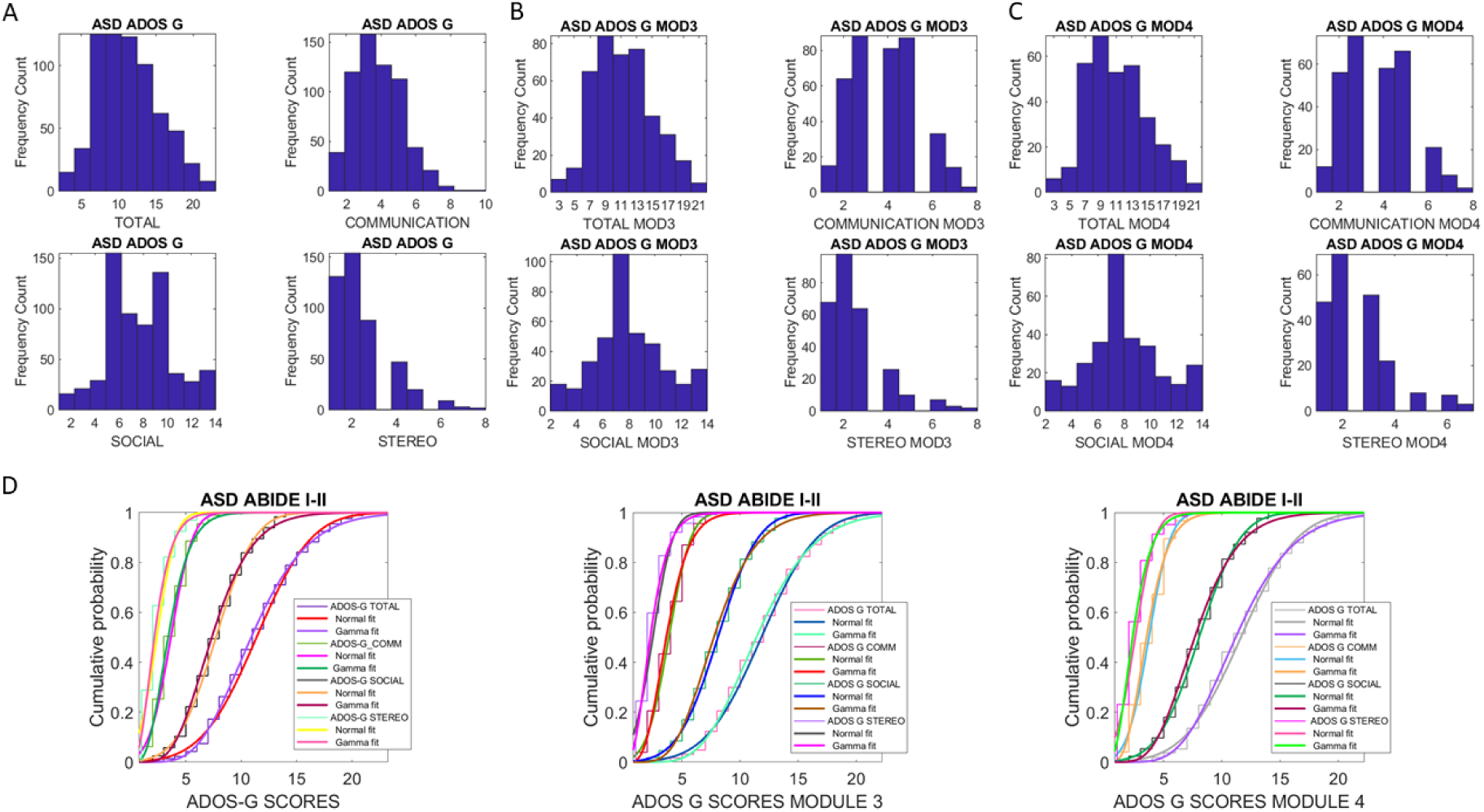
Non normality of the ADOS-G total and each of the sub-scores distributions in the ABIDE I, II sites. (A) Empirically obtained frequency histograms for each of the ADOS-G total and sub-scores across all ABIDE scores. (B) Same as in (A) but restricted to Module 3 scores. (C) Same as in (A) but restricted to Module 4 scores. (D) Empirically estimated CDFs for each sub-score for the Normal and Gamma distributions fit to the data. All cases failed the test of normality (see text) but no statistically significant differences were found between the empirical CDFs generated by Modules 3 and Module 4. Similar to ADOS-2 in Figure 4, no differences were found either between each of the Modules and the pooled data in (A).

The Lilliefors test failed the normality test with p << 0.01 for each sub-score of each ADOS test version. We also used maximum likelihood estimation (MLE) and fit several distributions to assess the best fit with 95% confidence. The MLE revealed the continuous Gamma family of PDFs had a better fit than the Normal PDF. The results from the Normal and Gamma fits are shown in Figures 4D and 5D for each of the corresponding sub-scores’ CDFs of the ADOS-2 and ADOS-G respectively.

### 3.2. ADOS scores Cannot distinguish between Females and Males

The ADOS score data from ABIBE were divided into the male and female participants to ascertain (1) if the distributions of the total and sub-scores were symmetric (test for normality) and (2) if they had statistically different overall scores comprising social, communication and stereotypical, repetitive motions that could distinguish the two phenotypes. The motivation for this comparison emerged from physiological (somatic-sensory-motor) metrics that can automatically distinguish males from females of the ABIDE repository [40; 41] and from distinction based on motor patterns derived from natural voluntary behaviors [26; 51]. These patterns can also automatically separate females/males with ASD from females/males with AS (from the subset of ABIDE with a DSM-IV classification) [41]. We reasoned that given that the ADOS tests are used to drive the science of Autism, i.e. to correlate physiological data with it; it may be important to ascertain whether the variability in the rater’s scoring from these versions of the ADOS, could also automatically distinguish the male from the female phenotype (much as the inherent variability in the physiological data does).

Figures 6AB and 7AB show the distributions of scores for males (6AB) and females (7AB). In all cases, the normality test failed according to the Lilliefors test (p<<10-4). Recall here that this test returns a test decision for the null hypothesis that the data comes from a distribution in the normal family, against the alternative that it does not come from such a distribution. In all cases the test rejected the null hypothesis at the 5% significance level. As with the pooled data in Figures 4A-5A and those from the breakdown into scores from Modules 3 or Module 4 in Figures 4B-C and 5B-C, we used MLE to evaluate the fit of different distributions, which we show in Figures 6AB center panel for the Normal and the Gamma family of distributions fit to the empirical CDFs of the total and sub-scores (left side ADOS-G and right side ADOS-2.)

**Figure 6.**
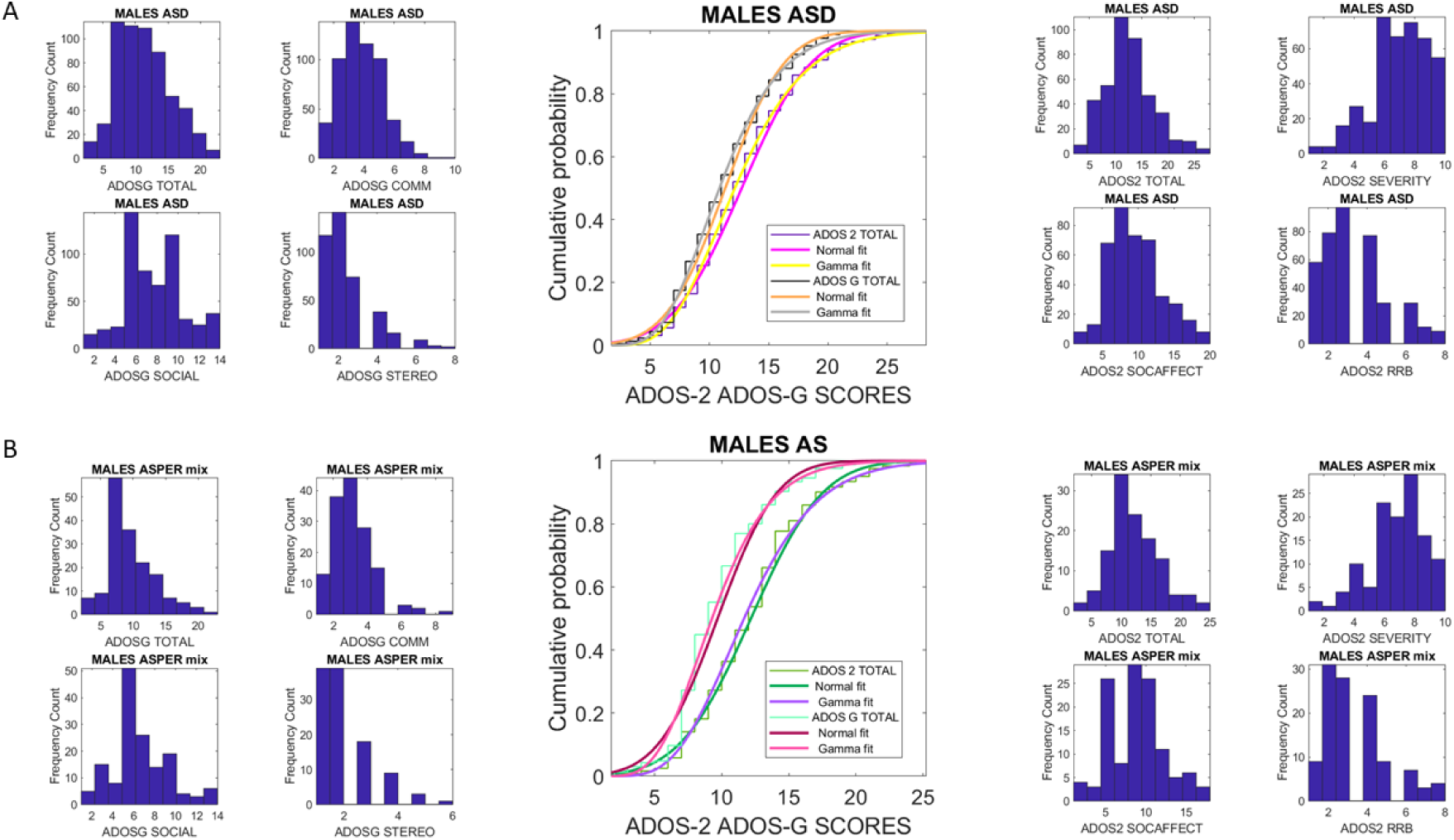
Non-normality of the distributions spanned by the males with ASD and those with Asperger’s syndrome according to the DSM-IV column of ABIDE. (A) Left panel shows the frequency histograms for each ADOS-G sub-score, while central panel shows the empirical CDFs fit by the theoretical Normal and Gamma family CDFs for the total score. Right panel shows similar plots for the ADOS-2. (B) Same format as in (A) for the case of Asperger’s syndrome (notice the separation of CDFs between ADOS-G and ADOS-2 contrasting with those of the ASD cases in (A)).

Despite the well-established physiological and neurobiological distinctions between males and females in the spectrum of Autism, here we could not find any statistically significant difference between the ADOS-G (or the ADOS-2) total scores to automatically separate these two distinct phenotypes. We could not find either statistically significant differences between the sub-scores of each ADOS test, when comparing males and females using the Wilcoxon ranksum test, as all *p-values* were above 0.5.

However, we did notice a significant difference in the distributions of the total scores for males when comparing those of the ADOS-G *vs*. those of the ADOS-2, (center panel in Figure 6B). This difference was assessed using the Kolmogorov-Smirnov test for 2 empirical distributions and yielded *p-value* of 0.0015. In contrast, this statistically significant difference was absent in the ADOS-G *vs*. ADOS-2 comparison of total scores from the females’ data with a *p-value* of 0.8 (Figure 7A center). Note here that these comparisons do not have meaningful clinical value. They are exclusively taken within a research framework to learn whether two versions of the same test always yield consistent separation; or to learn if in some cases they do not.

**Figure 7.**
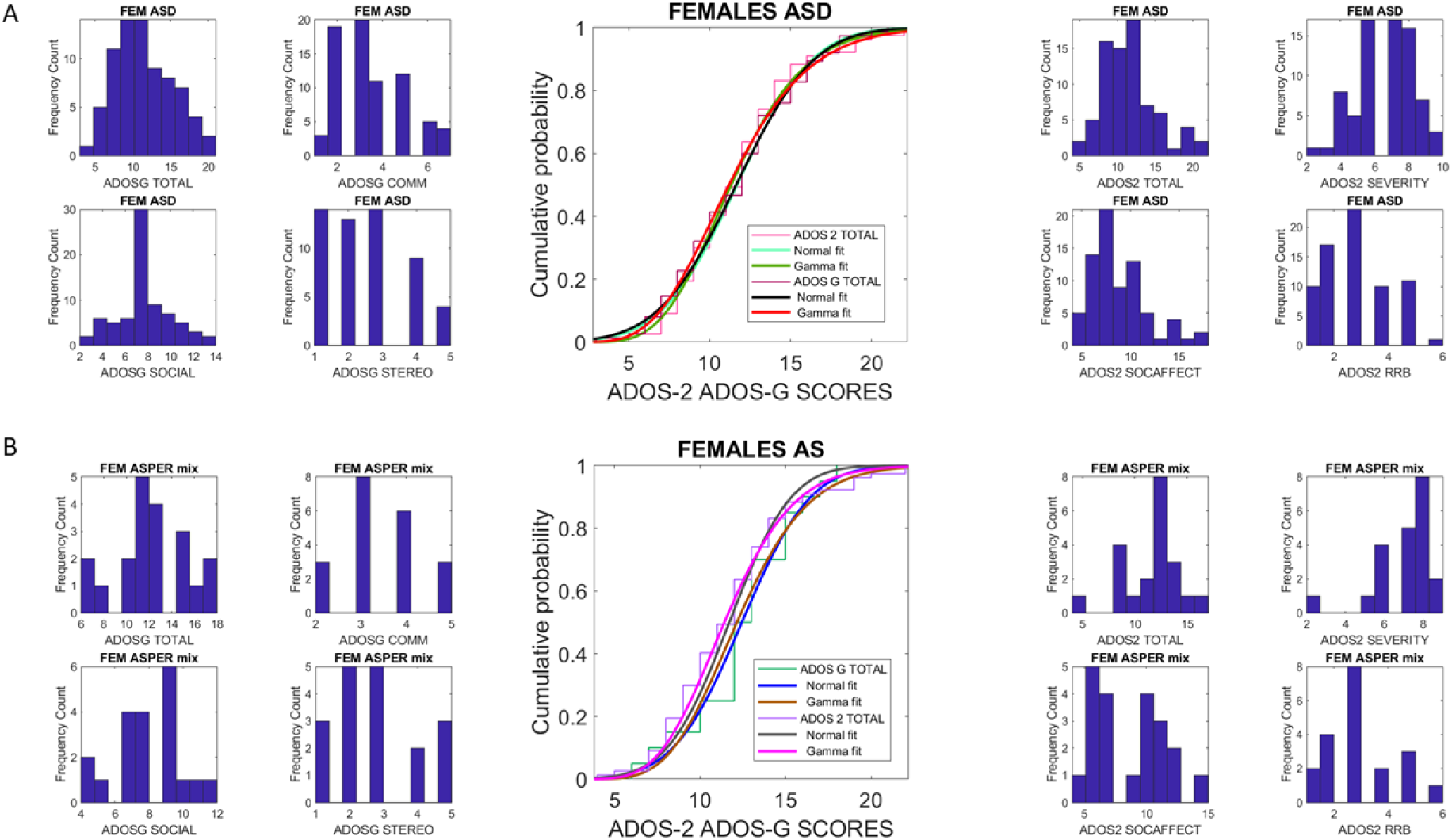
Non-normality spanned by the distributions of the females with ASD and AS in the ABIDE repository. (A) Frequency histograms of the ADOS-G total scores and sub-scores (left panel) and ADOS-2 (right panel) with center panel showing the empirical CDF fit by the Normal and Gamma distributions for the ADOS-G and ADOS-2 scores. (B) Same as in (A) for the females with a DSM-IV diagnosis of Asperger’s syndrome.

Given the visible separation of ADOS-G and ADOS-2 scores for the males, we proceeded to further interrogate the cohort of participants with AS according to the DSM-IV classification. This is possible as ABIDE provides the information on DSM-IV *vs*. DSM-5 on two separate columns of the data matrix. We then asked if the males with AS were also separable from the males with ASD, according to the ADOS scores from the two versions of the same test. Notice here the relevance of this question, as the ADOS-G and ADOS-2 are indistinctly used in Autism research (as instantiated by the ABIDE repository), and no differentiation is ever made by peer-reviewed papers that use one or the other to inform and guide the results from their physiology/neurobiology-based research.

### 3.3. Incongruent Results Between ADOS-G and ADOS-2 When Comparing Males with ASD and AS

We expected that despite their subtle differences, the variants of the same test would provide consistent results for participants with AS and for those with ASD. In the case of ADOS-G, we found a statistically significant difference between the scores of males with ASD and those with AS, using the Wilcoxon ranksum test yielding a *p-value* of 3.7×10^−7^. Further, the Kolmogorov-Smirnov test comparing 2 empirical distributions yielded a *p-value* of 2.4×10^−6^, thus rendering the two distributions for AS and ASD statistically different at the 0.05 level (i.e. rejecting the null hypothesis that the two sets came from a similar distribution.) In contrast to the ADOS-G, the ADOS-2 total score comparison (i.e. within the same test) between the males with ASD and AS, using the Wilcoxon ranksum test, yielded a *p-value* of 0.31, non-significant at the 0.05 level. The Kolmogorov-Smirnov test comparing 2 empirical distributions yielded a *p-value* of 0.5, with no significant difference and failing to reject the null hypothesis that the two data sets came from a similar distribution. Figure 6 shows the frequency histograms and CDF fits to the empirical data for males using the left-hand panel for the ADOS-G and the right-hand panel for the ADOS-2. Notice that the comparisons were made within each test, as they are different versions of the ADOS.

Comparisons of the females with ASD and AS are shown in Figure 7, using the total scores from the ADOS-G and ADOS-2 respectively. These outputs were congruent for the two versions of the test in that neither ADOS version distinguished the two groups. The Wilcoxon ranksum test yielded a *p-value* of 0.30 for ADOS-G and 0.54 for ADOS-2. The Kolmogorov-Smirnov test comparing 2 empirical distributions yielded a *p-value* of 0.42 for ADOS-G and 0.96 for ADOS-2. Notice that in both males and females breaking the data into Module 3 or Module 4 scores made no difference. As in the case of Figures 4 and 5, the distributions retained the shape and dispersion across the Modules, thus suggesting that blind classification of participants is not possible using these tests. In other words, given the scores of a Module 3 or a Module 4 version containing different tasks, it is not possible to know if they came from Module 3 or Module 4, as they span identical distributions. The numbers are statistically indistinguishable, so any machine learning algorithm attempting to classify participants based on these scores would fail, despite their coming from entirely different sets of tasks, and being aimed at assessing individuals with disparate language/communication levels.

We note that given the differences in sample size and the non-normality in the distributions of scores, in all above comparisons, we used the Wilcoxon rank sum test. This is a nonparametric test for two populations, used when samples are independent and of different sizes. It is equivalent to the non-parametric Mann-Whitney U-test for equality of population medians. The test statistic which ranksum returns is the rank sum of the first sample [52].

### 3.4. Normative Data Significantly Departs from Lowest-bound 0-score

The analyses of the in-person visit to assess the ADOS-2 in 52 participants, 26 suspected (and confirmed) to have Autism or ASD and 26 typical controls, yielded significantly different distributions of scores between the two age- and sex-matched groups. Figure 8A shows the frequency histograms of the ASD (top) and typical control (bottom) groups. Figure 8B shows the outcome of the non-parametric Kruskal-Wallis (one-way ANOVA) test highlighting the statistical significance of differences captured by the comparison. Further, Figure 8B rightmost panel highlights the output of this non-parametric test for the comparison of the 0-score denoting the absence of behavioral symptoms. For typical controls, the scores from the empirical data spanned a distribution of non-zero values, evident at the p<<.01 level.

**Figure 8.**
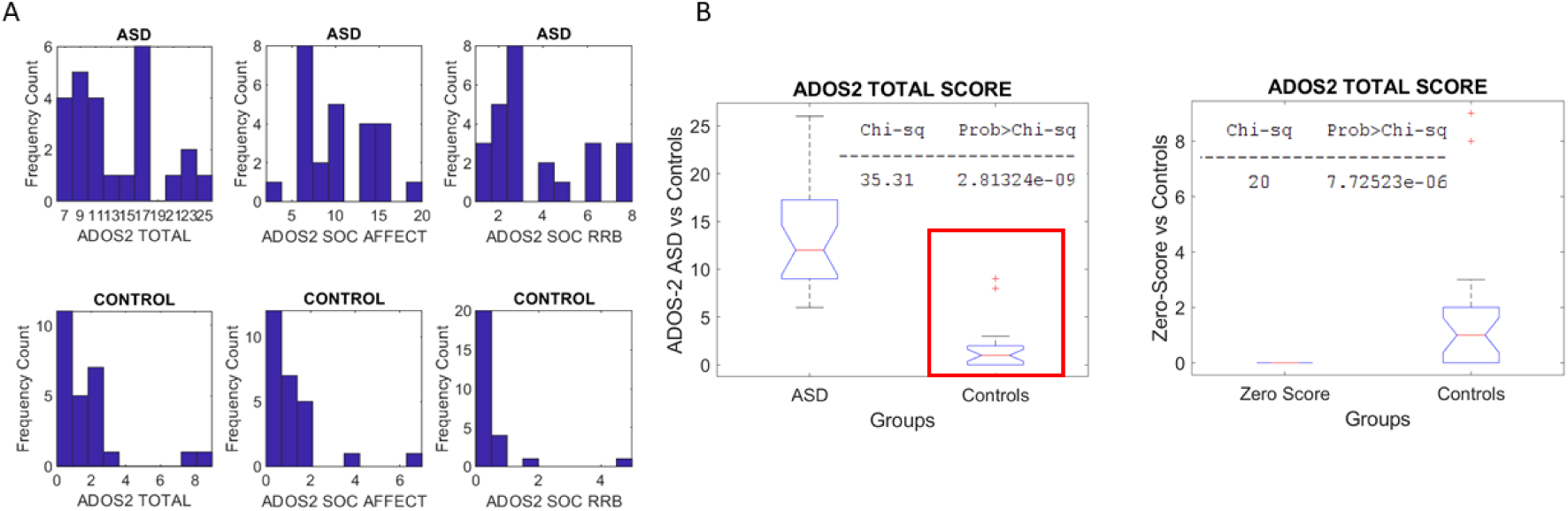
The need for normative data to build a true similarity metric for Autism research that adopts the ADOS test. (A) Frequency histograms of scores from the ADOS-2 rated scores for individuals with ASD and Autism cutoffs vs. those of typical controls. (B) Differences with statistical significance between the two groups total scores and (right panel) a distribution of scores for typical controls calling for a reassessment of the ADOS scoring system to convert it from a criterion-reference to a norm-reference system and build an appropriate metric space, amenable to measure change and derive developmental trajectories in basic scientific research.

**Figure 9.**
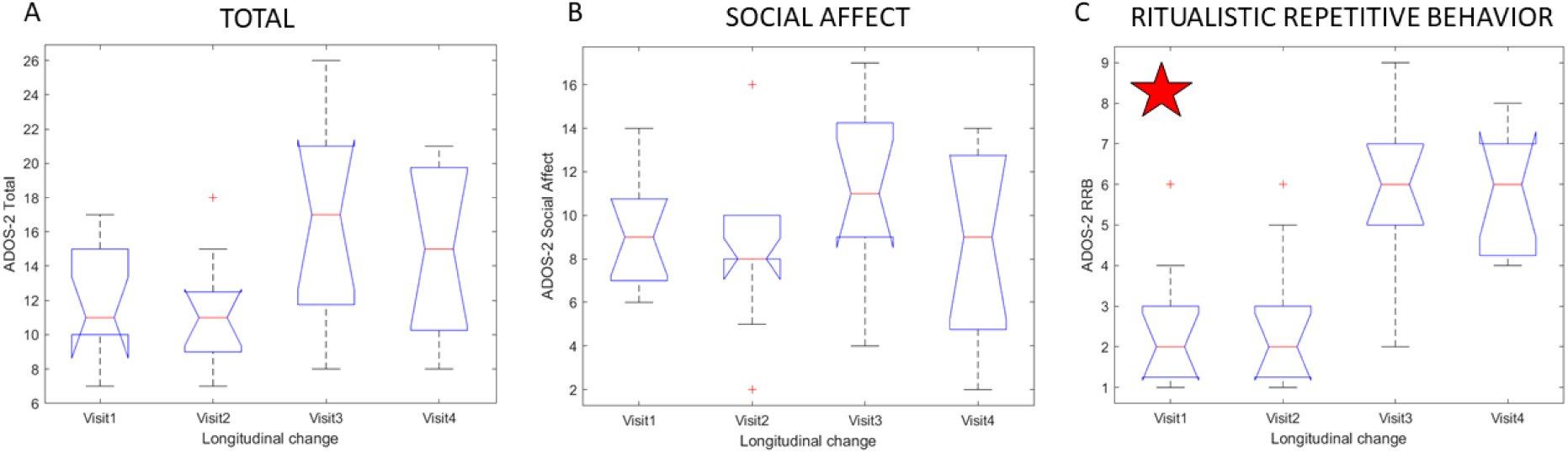
Susceptibility of RRB ADOS-2 sub-score to the rater clinician. (A) Output of the Kruskal-Wallis non-parametric one-way ANOVA using box and whiskers format show differences in total score between clinician 1 in visits 1-2 and clinician 2 in visits 3-4 but these differences do not reach statistical significance p<0.06. Notice the broader ranges in the score distribution of the clinician 2 and the presence of higher scores overall. (B) Social affect sub-scores were not significantly different between clinicians. (C) RRB scores were significantly different for the same children, same modules and same task order. Clinician 1 rated the children significantly lower with overall ranges contributing to a bias towards Autism spectrum. In contrast the higher values of the rates by clinician 2 contribute to a bias towards Autism diagnosis for the same children.

### 3.5. ADOS-2 RRB Outcome is Significantly Affected by Clinician

The longitudinal data across 4 visits enabled us to examine the influences of the clinician in 14 children that returned to the lab to perform the ADOS-2 test. The scores from all the children were pooled and the total score was compared across visits using the non-parametric Kruskal-Wallis test followed by the mult-compare test. The total score comparison revealed no significance (Chi-sq 7.21 and p-value 0.06.) Yet, given the borderline value close to 0.05 significance level, we examined the social affect score and the ritualistic repetitive behavior (RRB) score making up the total. We found no differences across visits in the social affect score. However, the RRB score significantly changed across visits (Chi-sq 21.01 and p-value 0.0001) with major differences when switching clinician in visit 3-4. Despite the use of the same modules, room setup and task order for each child, the differences in ADOS-2 scores for RRB were marked as systematically difference by the post hoc mult-compare test. These outcomes can be appreciated in Figure 10 for total (A), social affect (B) and RRB (C).

**Figure 10.**
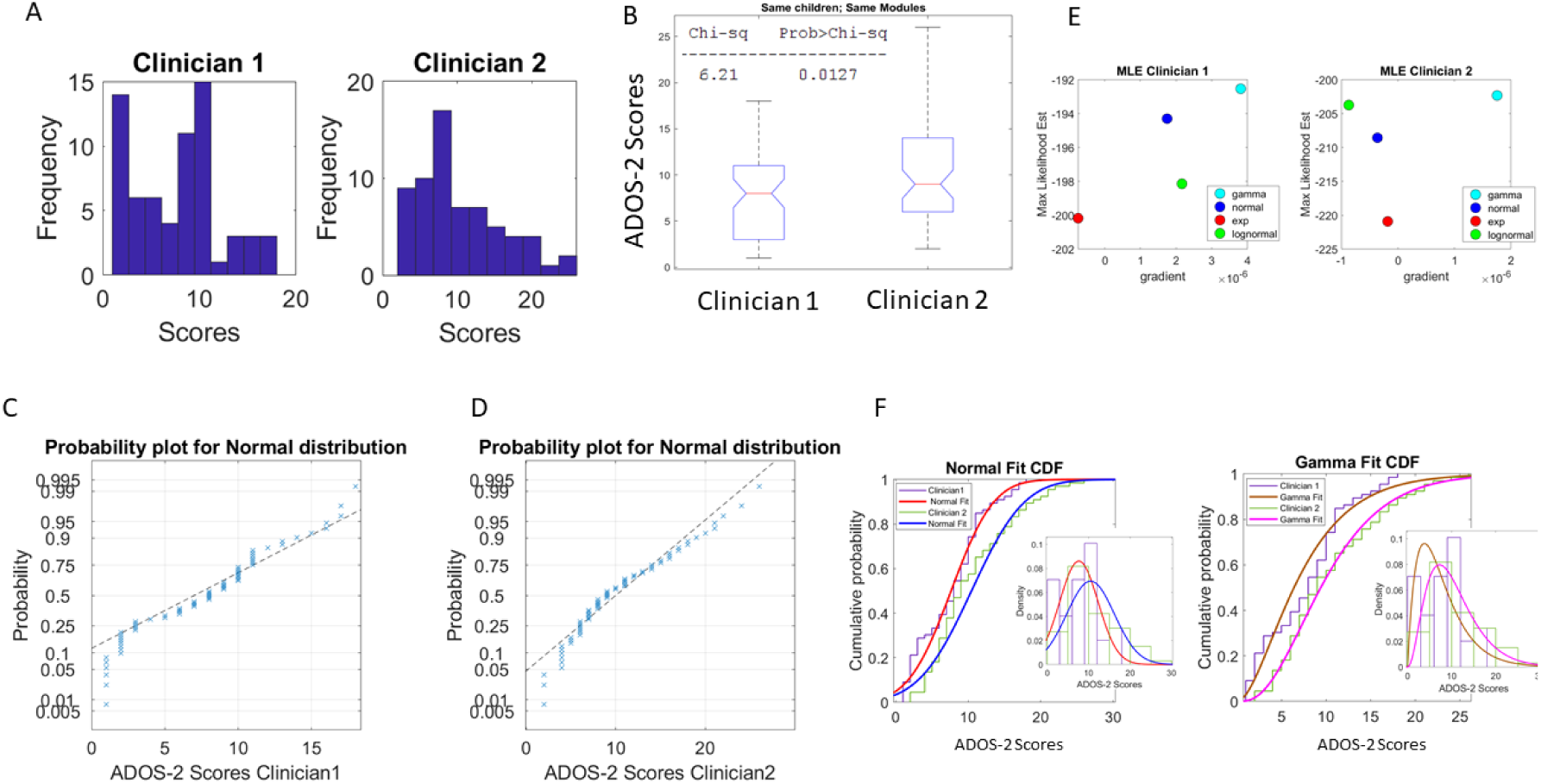
Non-normality of overall scores from each rater clinician. (A) Differences in rater clinician styles can be appreciated in the frequency histograms of the scores that each one obtained from the same cohort. (B) Differences in scores for the same cohort reach statistical significance (despite the relatively small number of measurements 14 children × 4 visits ×2 sub-scores, 112 scores). (C-D) Non-normality of the score samples from each rater clinician. (E) MLE output renders the normal distribution inappropriate in each rater clinician case. (F) Fitting the Normal vs. the Gamma family for each score set empirical CDF underscores the fundamentally different stochastic signatures derived from the variability of their rating of the same cohort of children with the same ADOS-2 modules.

We further tested all the scores for each clinician by pooling across all children and score type, to examine the types of distributions best fitting their frequency histograms. Figure 10A shows this analysis for each clinician, while Figure 10B shows the output of the non-parametric Kruskal-Wallis test, which revealed statistically significant difference. Figure 10C shows the failure of normality for scores by Clinician 1, while Figure 10D shows so for Clinician 2. The use of MLE to ascertain the fit of several probability distribution functions confirmed that the normal is not a good fit for either (Figure 10E). Further, the Gamma distribution was used as per the MLE outcome to fit the data and compare the scores of the two clinicians for the same children, same modules, same modules order/visit and same tasks order. Figure 10F shows the fits of the Normal distribution (left hand side) and that of the Gamma distribution (right hand side.) The Gamma distribution fit was best for Clinician 2 but poor for Clinician 1. The Normal was poor for both. The lognormal and exponential were also poor fits.

### 3.6. Age-corrected ADOS-2 Derivative Scores Confirm the RRB Score as the Most Affected by Change in Rater Clinician

The relative changes in score/age (with the age measured in years, months) were obtained for each of the 14 participants that we tracked over 4 visits. When examining these derivative data, we found significant differences in the RRB ADOS-2 scores. Consistent with the effect that the change of clinician in visit 3 had revealed for the size-data given by absolute ADOS-2 scores, here, the derivative-data considering the age change of the participant from visit to visit, also reveal significant changes in the RRB scores. The same individuals were rated significantly different by the clinicians, thus yielding different scores for the same module and tasks. Kruskal-Wallis test for the comparison of the ADOS-2 RRB scores across visits yielded significant differences across visits (Chi-sq 13.45, *p-value* 0.003).

The Figure 11A shows the evolution of the clinician’s diagnostic criteria over time. Visits 1 and 2 with clinician 1 show 10/14 with Autism vs. 4/14 with ASD. This pattern changes in Visit 2 for this rater clinician to 6/14 with Autism and 8/14 with ASD. The second clinician scores rather differently the same cohort of children performing the same tasks under the same room set up, from the same modules, compared to the rater clinician 1 in visits 1-2. In visits 3-4, the rater clinician 2 scores these same children as 50-50 Autism-ASD. The individual evolution for each child is seen in Figure 11B, whereby the different clinician’s styles of scoring can be seen. There is no inter-rater reliability rendering these ADOS-2 criteria robust. For the same child and ADOS-2 module, we see changes in the classification of Autism vs. ASD. These differences in perception biases add to the findings on non-normality of the distributions previously described in Figure 10 for this lab cohort, and for the large cross-sectional population data from ABIDE.

**Figure 11.**
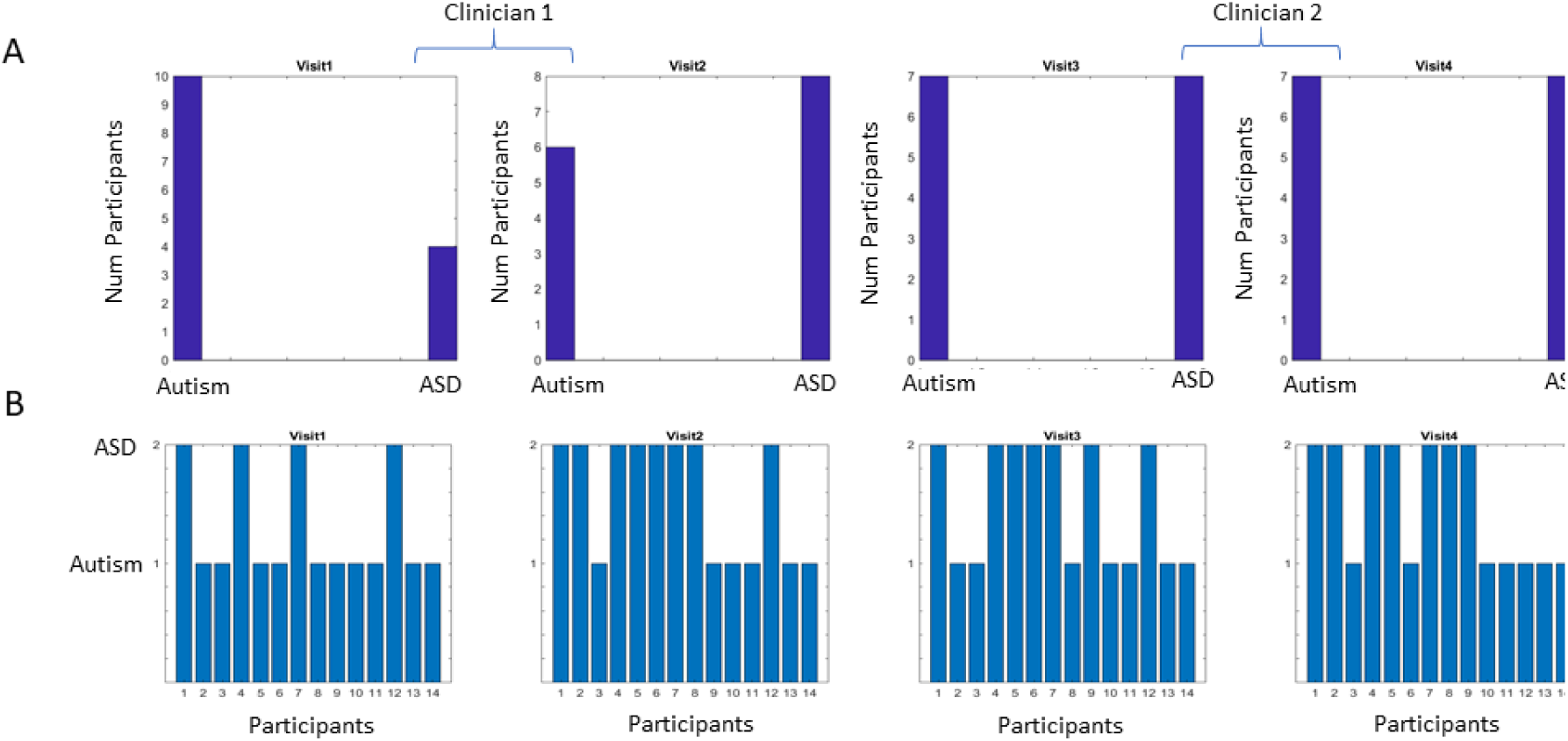
Non-reliable diagnostics for the same cohort of children under the same ADOS-2 modules and tasks. (A) Bias of rater clinician 1 vs. rater clinician 2 reveals different styles bound to impact the outcome. (B) Variability of diagnostic classification (Autism vs. Autism Spectrum) for each of the 14 participating children in the four visits spanning 1.3 years on average ± 6 months.

## 4. Discussion

This work examined the statistical signatures of the variability inherently present in research-grade ADOS-G and ADOS-2 scores adopted by researchers and available in the ABIDE repository. With 1,324 full clinical records, and a smaller pilot sample of 52 participants (26 diagnosed in the laboratory and 26 typical controls) the work set to examine the statistical requirements affecting the relationship between bias and sensitivity. The purpose of the study was several fold: (1) test the assumption of normality inherent in the current *one size fits all* clinical model of Autism used in research and informing and largely steering basic scientific research; (2) highlight the need for a norm-reference clinical metric leading to a proper similarity metric to measure change and its rate in neurodevelopment; (3) test the inter-rater reliability by characterizing the rater’s bias under similar conditions (i.e. same cohort of children performing the same two modules and tasks in the same laboratory-controlled environment) and (4) identify which sub-score would be the most affected by the change in rater-clinician.

The results from the analyses revealed that the research-grade ADOS scores reported by researchers in ABIDE generate non-normal distributions that change shape and (scale) dispersion for the total score and across all the sub-scores, for each of the ADOS-G and ADOS-2 tests. The scores confound males and females (unlike the ABIDE physiological data from the exact same participants, which distinguishes males *vs.* females and ASD *vs.* AS [5; 40; 41]). More alarmingly, the ADOS-G and ADOS-2 revealed fundamentally different diagnoses for AS individuals under DSM-IV criteria. Furthermore, when these AS scores were compared to those of ASD under the ADOS-G, the results were very different than when they were assessed with the ADOS-2. This is highly problematic, given that the research-grade ADOS is systematically used in the scientific literature, without ever distinguishing between the ADOS-G or the ADOS-2 versions.

These versions of the instrument have indeed been used in basic scientific research papers, but perhaps some points should be considered in future studies:

- The empirical distributions of these scores are not normal.
- There is no proper metric space to build a similarity measure that enables us to assess departure from typical development over the human lifespan, *in an age-dependent manner*.
- Because there is no similarity metric for this test (as the ADOS is not a norm-referenced test), repeating the test overtime and using it to describe neurodevelopmental ***change*** and build neurodevelopmental trajectories in autism is mathematically inappropriate and misleading.

The most important message of this work is that without a proper metric scale, it will not be possible to capture change in neurodevelopment. Typical neurodevelopment occurs at accelerated rates of change with non-normal distributions across parameters of physical growth and neuromotor control [50]. Neurodevelopment in autism occurs at non-linear, non-uniform rates whereby age does not have the same meaning as it does in neurotypical development [6].

The nervous system of an individual with autism is a coping system with self-correcting mechanisms that evolve over time and give rise to non-uniform developmental trajectories [39]. This means that a transformation-operation that e.g. considers age, to change non-normal distributions into normal distributions (i.e. to call it a standard measure as in the standardized severity score) will not address the fundamental problem of the individual that is developing with a coping nervous system. The problem that we bring for consideration to the scientific community doing research on autism is not one of the statistical (in)appropriateness of a test, but rather of the need for tests that reflect the self-correcting nature of nervous systems. Instead of masking the phenomena of relevance with some transformation of the data to make up for normality, or altogether enforcing normality, we need to use a data-driven approach.

Data-driven approaches tend to preserve empirically assessed features of phenomena. They do not throw away important variability and as such, offer the possibility of capturing the true nature of ***change*** in a coping neurobiological system that is developing at atypical rates. Although this paper uses the ADOS test as the example to illustrate the potential problems that blindly adopting such tests for scientific research may create, the same tenets apply to any other clinical test used in basic research of neurodevelopmental disorders. These disorders reflect in great part problems with the nervous systems and since nervous systems are adaptable, we would be missing self-correcting mechanisms by imposing theoretical models without empirically informing those models.

In the context of this test, changes in the distributions’ shape and dispersion (across ages and sex) imply lack of independence between sensitivity and bias. As such, the rater’s inherent bias and the non-normality of scores’ probability distributions cast doubt on the appropriateness of signal detection theory (SDT) and the Receiver Operating Characteristic ROC-area-under-the-curve as criteria testing the sensitivity and reliability of this test.

There remains an absolute lack of a proper standard range of scores with physiological interpretation, despite claims that the ADOS has been standardized [35; 36; 43; 53; 54; 55].

Given these findings, it may be important to reconsider adopting this test to inform basic scientific research in neurodevelopment. Across a multitude of research papers, discrete scores that do not have a proper metric are systematically forced to be (linearly) correlated with continuous physical data, yet the lack of normality in the distributions of the scores along with the lack of independence between raters’ bias and sensitivity pose a problem for its validity in research, according to SDT and ROC area-under-the-curve types of analyses [56]. One now wonders how many false positives we may have in research studies. What does it really mean to have Autism, or to be on the Autism spectrum? And how can that distinction be made relative to normative data from typically developing controls, when no such data exist? Here we see, even in a rather modest cohort of neurotypical participants, that there is not such a thing as a 0-score (or even epsilon-value score) for typical neurodevelopment, thus indicating the presence of behavioral symptoms in the neurotypical population. Given such fluctuations, how can we build a proper metric for neurodevelopmental research?

As stated, neurodevelopment occurs at highly non-linear accelerated rates. In a coping nervous system, age is a non-uniform quantity in that any two given children with the same age may have very different levels of nervous systems maturation [6; 39] (Figure 1). Despite the lack of metric and non-normality of the ADOS scores, there have been recent longitudinal studies that have used the ADOS scores to attempt to measure developmental trajectories by repeating the test on the same participant, over time. This has been done by taking the (assumed) theoretical Gaussian mean across scores, while repeating the process and expressing the plots of highly non-linear developmental data on linear scales [21]. Indeed, such statistically problematic work has been rendered by lead researchers in autism as new standard to map early symptom trajectories in ASD [20]. This lack of basic statistical appropriateness poses a challenge for the rest of the community as it produces misleading inferences and interpretations of such longitudinal data.

Open access repositories now make it possible to examine (for the first time by researchers from different fields with different skillsets) the validity of the use of this instrument in research, using unprecedented large number of records. These provide enough statistical power, where the high cost of running these studies often prevents labs from deploying them. For example, recent work using the National Database for Autism Research (NDAR) and the Simons Simplex collection demonstrated that it is possible to shorten the administration time of the ADOS, while preserving cutoff criteria. While this earlier work already highlighted the non-normality of the distributions of the scores reported in those data repositories [57], the present work takes a deeper look at the assumptions that the research-grade test makes. Here we further raise several relevant questions about the need for normative assessment across the human lifespan, to truly formulate a standard test with a proper metric for research use in neurodevelopment.

Perhaps the use of the ADOS in the clinical arena is less damming than the use of its research-grade version in basic scientific research. In the US, the Autism label ensures coverage for certain treatments that many children could not otherwise have access to (particularly in under-represented minorities and poor areas.) Even though the ADOS testing does not produce an official diagnosis, its use in research labs in the US, could serve as a flag to send parents to federally certified clinics that offer services upon multi-prone criteria involving other tests. However, owing to copyright issues, the Western Psychological Services Firm, WPS company does not allow researchers copy, reproduce or share the ADOS booklet with important details of the outcome. In other words, those children that come to our labs and receive the research-grade ADOS and pass the cut-off scores are labeled autistic by this test. However, as researchers, we are not allowed to share details with their parents. This obstructs their ability to go to a proper clinic and pursue the diagnosis that will give them access to Early Intervention Programs or to Individualized Education Programs in cases when the child is of school age. If the WPS and the trainers of the ADOS allowed this, the test adopted by researchers would serve as a warning to parents that some aspects of the child’s neurodevelopment may be off track.

Unlike the statistical confounds that the ADOS total scores and sub-scores surely bring to research, in its current form at the clinic, the ADOS provides psychological comfort to adults who had never been previously diagnosed and could not understand their place in the social scene. Many adults, newly diagnosed at the clinic, express a sense of relief upon learning that they are on the Autism spectrum and as such have social-interaction differences. Further, the ADOS adds important information to the coarser DSM diagnosis. Thus, the clinical value of this instrument is highly appreciated. However, its use as a research instrument to inform physiological studies, is clearly now questionable and should be discussed among the scientific community, particularly the community with the skill set to fully understand statistics beyond black-box approaches that utilize software packages, i.e. without ever verifying the assumptions of the methods implemented by those packages.

We should point out that the SDT framework used by the ADOS to test validity, was introduced to science in the late 50’s to address very different problems in engineering. It was adapted to Psychophysics in Psychology under very different conditions than the types of social interactions presumably tested by the ADOS would require. Here we have a test of social interactions, i.e. a physical dance that takes place as a dialogue between two human beings. Yet, this test is treated as a monologue. In this monologue, the same person that sends the message, receives it and scores it, thus acting simultaneously as the stimulus and the response, plus some ‘noise’ produced by the variability in the responses by the individual being observed, assessed and rated. Although the test provides a protocol for structured social exchange, in its present form, this test cannot tell us much about the types of somatic-sensory-motor issues that are by now well-established in Autism by the scientific community [4; 6; 27; 58; 59; 60; 61; 62; 63; 64; 65; 66; 67; 68; 69; 70; 71] and recognized by the inclusion of sensory criteria in the DSM-5. For example, the most problematic sub-score in this example, the RRB, is inherently a sensory-motor component difficult to predict, owing this to its somewhat random appearances. Much like predicting a seizure or anticipating a self-injurious episode, RRBs spontaneously emerge in inexplicable forms, sometimes seemingly independent of the rater’s abilities or training, sometimes seemingly triggered by the uncertainty that the ADOS questioning brings to the person under examination. To predict them with high certainty, we would have to first characterize them with instruments that pickup information that escapes the naked eye.

There is room for transformative change bound to improve the use of such clinical tests in scientific research studies in general. After years without recognizing the importance of sensory motor issues in mental health [72], the Research Domain Criteria (RDoC) matrix created by the NIMH [11; 73] has finally included in January of 2019 the entry for sensory-motor issues. This new development, paired with the admission by the DSM-5 that there are sensory issues in Autism, will provide a new foundation to explore human social behaviors from a new angle that can include movements and their sensation [14], while offering the opportunity to uncover the inherent capabilities and predispositions that a coping nervous system develops [39].

The ADOS test nevertheless still fails to recognize sensory-motor issues [1], as we quote the following caveat when choosing a module from the manual: “*Note that the ADOS-2 was developed for and standardized using populations of children and adults* ***without significant sensory and motor impairments.*** *Standardized use of any ADOS-2 module presumes that the individual can walk independently and is free of visual or hearing impairments that could potentially interfere with use of the materials or participation in specific tasks*” (emphasis added) Catherine Lord, Rutter, DiLavore, Risi, & Western Psychological Services Firm.))”.

Interestingly, despite this caveat for the use of the ADOS test, stated in their manual, to the best of our knowledge, the makers of the ADOS test have never reported scientific studies of individuals with autism where it is objectively established that individuals in the spectrum of autism have no significant sensory and motor impairments. Yet, invariably, when we test the children in basic scientific research labs, using high grade instruments, we do find visual, hearing and touch impairments that would surely interfere with the use of the materials in this test. These highly quantifiable problems with their somatic and sensory motor systems are the tip of the iceberg, as deeper problems are present with their enteric nervous systems (the gut) and microbiome [74; 75; 76; 77; 78]. Many suffer from pain and temperature dysregulation as their overall sense of touch, vestibular issues with balance and multi-sensory integration overwhelms them in ways that we can now precisely quantify in personalized manner. Perhaps the new NIH-RDoC sensory-motor criteria will help WPS redefine ADOS for research and encourage the use of new objective criteria grounded on biophysical metrics assessing the nervous systems’ functions.

There is by now mounting evidence that somatic-sensory-motor issues do exist across the many phenotypes that go on to receive this diagnosis today under the DSM-5 broader criteria. The DSM-5 allows Attention Deficit Hyperactivity Disorder (ADHD) and sensory issues in the criteria for Autism. As such, Autism is no longer a narrowly, well-defined disorder (perhaps it never was) despite insistence on defining it by clinical criteria that does not reflect the underlying physiological conditions that these individuals possess from birth onward. A new emergent field aimed at the uncovering of multiple digital biomarkers to characterize and automatically stratify various aspects of behavior for research purposes, may be the answer to the start of a new era in Autism scientific research aimed at a physiological characterization for medical use. In our present study, it was evident that whether using absolute scores, or derivative, age-dependent data accounting for longitudinal dynamic changes from visit to visit, the RRB reflecting sensory motor issues, picked up best the switching of the clinician. If we were to combine this structured social test with wearable biosensors, we could automatically stratify Autism spectrum disorders and provide objective criteria of use to the community doing basic scientific work (e.g. geneticists, electrophysiologists, neuroimaging, etc.) and to the physicians treating the medical issues. Some collaborative work along those lines has been done between clinicians and researchers, but more research is needed to fully validate and replicate the use of digital ADOS within the smart-mobile and personalized health concepts.

## 5. Conclusions

We invite the readership to consider that science in Autism needs to retake the path of independence and reclaim its agency to be able to conduct proper scientific research. This can be done by building an inter-disciplinary consortium of scientists from diverse disciplines with the skill set to derive proper metrics and true standardized methods for personalized medicine. The misinformation in some cases, or lack of information in other instances, that the scientific community has been subject to can easily stop by taking advantage of open access data sets and teaming up with experts in the more exact sciences. More generally, across all disciplines, we need to empirically verify inherent assumptions and understand in the first place what those assumptions mean in the models that we use. Peer review within an insular community will not work in favor of scientific progress. Autism is much too important to ignore these recommendations. It is the future of millions of children that we scientists have in our hands. *Should we disrupt this lack of rigor and transform Autism science by informing it with objective means?*

## Supporting information

Graphical Abstract

## Author Contributions

conceptualization, E.B.T.; methodology, E.B.T.; software, E.B.T., S.M.; validation, E.B.T., R.R., S.M. and B.G.; formal analysis, E.B.Y.; investigation, E.B.T., R.R., S.M., B.G.; resources, E.B.T.; data curation, E.B.T., R.R., S.M., B.G.; writing— original draft preparation, E.B.T.; writing—review and editing, R.R., S.M., B.G.; visualization, E.B.T.; supervision, E.B.T.; project administration, E.B.T.; funding acquisition, E.B.T.

## Acknowledgments

We thank the children and families who participated in this study. We thank the anonymous clinicians who helped us with administration of the ADOS, scoring and reliability assessment. We thank the past and current members of the Rutgers Sensory Motor Integration Lab who helped with data collection and study coordination.

