## Supplemental Table for "Hidden Aspects of the Research-ADOS are Bound to Affect Autism Science"

**Table 1** Baseline ADOS-2 with 52 participants, 26 participants with ASD and 26 controls

| ID ASD | Age | V1 Mod | V1 SA | V1 RRB | V1 Total | ID Ctrl | Age | V1 Mod | V1 SA | V1 RRB | V1 Total |
|---|---|---|---|---|---|---|---|---|---|---|---|
| 1 | 4 | 3 | 7 | 3 | 10 | 1 | 8 | 3 | 0 | 0 | 0 |
| 2 | 8 | 3 | 6 | 3 | 9 | 2 | 10 | 3 | 0 | 0 | 0 |
| 3 | 10 | 3 | 14 | 3 | 17 | 3 | 9 | 3 | 1 | 1 | 2 |
| 4 | 13 | 3 | 6 | 1 | 7 | 4 | 12 | 3 | 2 | 0 | 2 |
| 5 | 6 | 3 | 6 | 3 | 9 | 5 | 7 | 3 | 0 | 0 | 0 |
| 6 | 6 | 3 | 6 | 3 | 9 | 6 | 7 | 3 | 2 | 0 | 2 |
| 7 | 11 | 3 | 10 | 2 | 12 | 7 | 10 | 3 | 4 | 5 | 9 |
| 8 | 5 | 1 | 15 | 3 | 18 | 8 | 9 | 3 | 0 | 0 | 0 |
| 9 | 9 | 1 | 11 | 2 | 13 | 9 | 7 | 3 | 1 | 0 | 1 |
| 10 | 6 | 3 | 14 | 3 | 17 | 10 | 11 | 3 | 1 | 0 | 1 |
| 11 | 14 | 1 | 7 | 1 | 8 | 11 | 7 | 3 | 2 | 0 | 2 |
| 12 | 10 | 1 | 11 | 6 | 17 | 12 | 15 | 4 | 1 | 1 | 2 |
| 13 | 4 | 3 | 9 | 2 | 11 | 13 | 11 | 4 | 1 | 0 | 1 |
| 14 | 11 | 1 | 13 | 3 | 16 | 14 | 13 | 4 | 0 | 0 | 0 |
| 15 | 9 | 3 | 6 | 4 | 10 | 15 | 31 | 4 | 1 | 0 | 1 |
| 16 | 7 | 3 | 6 | 2 | 8 | 16 | 49 | 4 | 2 | 0 | 2 |
| 17 | 7 | 3 | 9 | 2 | 11 | 17 | 48 | 4 | 0 | 0 | 0 |
| 18 | 11 | 3 | 10 | 1 | 11 | 18 | 38 | 4 | 0 | 0 | 0 |

| | | | | | | | | | | | |
|---|---|---|---|---|---|---|---|---|---|---|---|
| 19 | 4 | 1 | 20 | 6 | 26 | 19 | 29 | 4 | 0 | 0 | 0 |
| 20 | 8 | 3 | 13 | 5 | 18 | 20 | 30 | 4 | 0 | 0 | 0 |
| 21 | 10 | 1 | 16 | 8 | 24 | 21 | 32 | 4 | 0 | 1 | 1 |
| 22 | 13 | 1 | 11 | 8 | 17 | 22 | 22 | 4 | 1 | 2 | 3 |
| 23 | 10 | 2 | 15 | 8 | 23 | 23 | 48 | 4 | 0 | 0 | 0 |
| 24 | 4 | 2 | 2 | 4 | 6 | 24 | 20 | 4 | 7 | 1 | 8 |
| 25 | 18 | 3 | 15 | 6 | 21 | 25 | 66 | 4 | 0 | 0 | 0 |
| 26 | 20 | 4 | 6 | 1 | 7 | 26 | 43 | 4 | 2 | 0 | 2 |

**Table 2** Longitudinal assessment of 14 participants across 4 visits, 2 different ADOS-2 modules and 2 raters.

| ID | Age | V1 Mod | V1 SA | V1 RRB | V1 Tot | V1 Dx | V2 Mod | Age | V2 SA | V2 RRB | V2 Tot | V2 Dx | V3 Mod | Age | V3 SA | V3 RRB | V3 Tot | V3 Dx | V4 Mod | Age | V4 SA | V4 RRB | V4 Tot | V4 Dx |
|---|---|---|---|---|---|---|---|---|---|---|---|---|---|---|---|---|---|---|---|---|---|---|---|---|
| 1 | 4.3 | 3 | 7 | 3 | 10 | **S** | 2 | 5.2 | 5 | 2 | 7 | **S** | 3 | 5.9 | 4 | 5 | 9 | **S** | 2 | 7.3 | 4 | 4 | 8 | **S** |
| 2 | 8.9 | 3 | 6 | 3 | 9 | **Aut** | 2 | 9.2 | 8 | 0 | 8 | **S** | 3 | 10.5 | 7 | 4 | 11 | **Aut** | 2 | 10.8 | 8 | 1 | 8 | **S** |
| 3 | 10.1 | 3 | 14 | 3 | 17 | **Aut** | 2 | 10.4 | 10 | 3 | 13 | **Aut** | 3 | 10.9 | 16 | 8 | 24 | **Aut** | 2 | 11.3 | 14 | 7 | 21 | **Aut** |
| 4 | 12.5 | 3 | 6 | 1 | 7 | **S** | 4 | 13.9 | 2 | 6 | 8 | **S** | 3 | 14.1 | 9 | 2 | 11 | **S** | 4 | 14.6 | 2 | 6 | 8 | **S** |
| 5 | 6.6 | 3 | 6 | 3 | 9 | **Aut** | 2 | 6.8 | 6 | 1 | 7 | **S** | 3 | 7.2 | 5 | 3 | 8 | **S** | 2 | 7.4 | 4 | 3 | 7 | **S** |
| 6 | 12.1 | 3 | 10 | 2 | 12 | **Aut** | 2 | 12.5 | 8 | 1 | 9 | **S** | 3 | 12.1 | 12 | 6 | 18 | **S** | 2 | 13.2 | 8 | 5 | 13 | **Aut** |
| 7 | 14 | 1 | 7 | 1 | 8 | **S** | 2 | 14.3 | 8 | 2 | 10 | **S** | 1 | 14.7 | 9 | 5 | 14 | **S** | 2 | 15.1 | 9 | 6 | 15 | **S** |
| 8 | 10 | 1 | 11 | 6 | 17 | **Aut** | 1 | 10.4 | 10 | 5 | 15 | **S** | 1 | 10.7 | 17 | 9 | 26 | **Aut** | 1 | 11.3 | 14 | 7 | 21 | **S** |
| 9 | 4 | 3 | 9 | 2 | 11 | **Aut** | 2 | 4.5 | 8 | 3 | 11 | **Aut** | 3 | 5 | 15 | 7 | 22 | **S** | 2 | 5.4 | 13 | 7 | 20 | **S** |
| 10 | 11.6 | 1 | 13 | 3 | 16 | **Aut** | 1 | 11.7 | 16 | 2 | 18 | **Aut** | 1 | 12.1 | 11 | 7 | 18 | **Aut** | 1 | 12.4 | 11 | 8 | 19 | **Aut** |
| 11 | 9.2 | 3 | 6 | 4 | 10 | **Aut** | 2 | 9.6 | 10 | 1 | 11 | **Aut** | 3 | 9.1 | 10 | 6 | 16 | **Aut** | 2 | 10.2 | 4 | 6 | 10 | **Aut** |
| 12 | 8.2 | 3 | 6 | 2 | 8 | **S** | 2 | 8.7 | 8 | 0 | 8 | **S** | 3 | 9.2 | 6 | 2 | 8 | **S** | 2 | 9.8 | 7 | 3 | 10 | **Aut** |
| 13 | 7 | 3 | 9 | 2 | 11 | **Aut** | 2 | 7.4 | 9 | 2 | 11 | **Aut** | 3 | 7.9 | 11 | 6 | 17 | **Aut** | 2 | 8.1 | 12 | 4 | 16 | **Aut** |

| | | | | | | | | | | | | | | | | | | | | | | | | |
|---|---|---|---|---|---|---|---|---|---|---|---|---|---|---|---|---|---|---|---|---|---|---|---|---|
| 14 | 11.9 | 3 | 10 | 1 | 11 | **Aut** | 2 | 12 | 8 | 1 | 9 | **Aut** | 3 | 12.4 | 6 | 2 | 8 | **Aut** | 2 | 12.1 | 7 | 4 | 11 | **Aut** |
