## Supplementary figures and images for "Hidden Aspects of the Research-ADOS are Bound to Affect Autism Science"

### Graphical Abstract

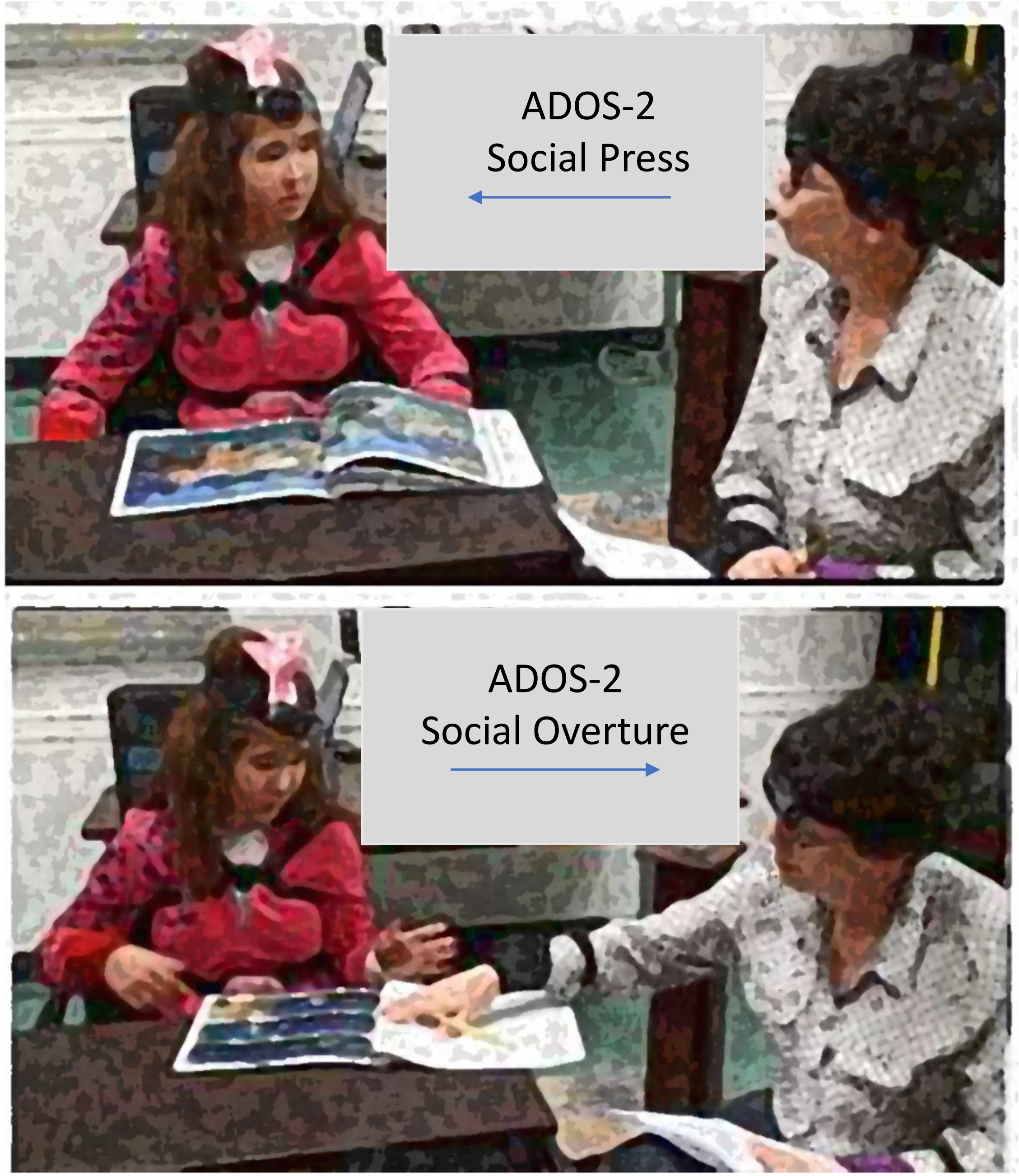

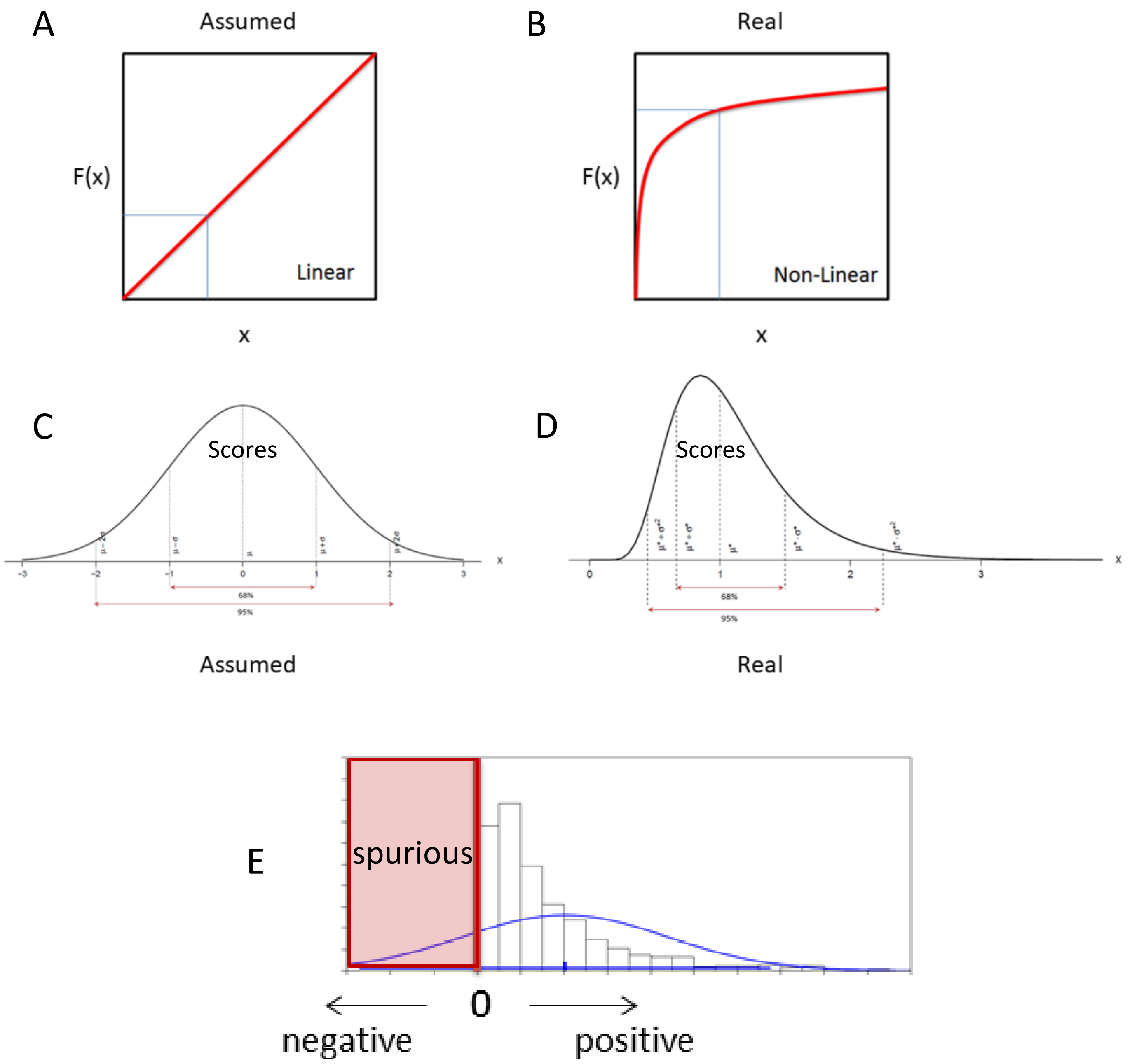
